# Germ fate determinants protect germ precursor cell division by restricting septin and anillin levels at the division plane

**DOI:** 10.1101/2023.11.17.566773

**Authors:** Caroline Q. Connors, Michael S. Mauro, J. Tristian Wiles, Andrew D. Countryman, Sophia L. Martin, Benjamin Lacroix, Mimi Shirasu-Hiza, Julien Dumont, Karen E. Kasza, Timothy R. Davies, Julie C. Canman

## Abstract

Animal cell cytokinesis, or the physical division of one cell into two, is thought to be driven by constriction of an actomyosin contractile ring at the division plane. The mechanisms underlying cell type-specific differences in cytokinesis remain unknown. Germ cells are totipotent cells that pass genetic information to the next generation. Previously, using *formin^cyk-1^(ts)* mutant *C. elegans* embryos, we found that the P2 germ precursor cell is protected from cytokinesis failure and can divide without detectable F-actin at the division plane. Here, we identified two canonical germ fate determinants required for P2-specific cytokinetic protection: PIE-1 and POS-1. Neither has been implicated previously in cytokinesis. These germ fate determinants protect P2 cytokinesis by reducing the accumulation of septin^UNC-59^ and anillin^ANI-1^ at the division plane, which here act as negative regulators of cytokinesis. These findings may provide insight into cytokinetic regulation in other cell types, especially in stem cells with high potency.

## Introduction

Germ cells play a unique role in passing genetic information from one generation to the next. Perhaps because germ cell integrity is critical for fitness and continuation of the species, there appear to be specific mechanisms to protect germ cell fate, survival, and proliferation. For example, germ precursor cells undergo specific differential developmental pathways.^1-10^ There is also evidence in several model systems that cytokinesis in germ precursor cells is differentially regulated from in somatic cells. Cytokinesis is the physical division of one cell into two, which occurs at the end of the cell cycle. In many metazoan germ cells, unlike somatic cell divisions^11^, daughter cells are not severed via abscission and remain connected by a stable intercellular bridge (for review see^12^). In *Drosophila*, mutations in the cytoskeletal interacting protein anillin specifically affects somatic cell but not germ precursor cell cellularization.^13,14^ In our own previous work, we found that cytokinesis in *C. elegans* germ precursor cells is uniquely resistant to severe perturbations of the actin cytoskeleton that completely block cytokinesis in somatic cells^15^ (see also^16^). Thus, cell division in germ precursor cells appears to have significant differences in regulation from somatic precursor cell division.

These data contradict the textbook view of cytokinesis that all animal cells divide using the same molecular machinery. It is thought that anaphase onset drives mitotic spindle signaling to promote the assembly and constriction of an actomyosin contractile ring at the cell division plane to power cytokinesis. In fact, growing evidence supports both cell type-specific regulation of cytokinesis and cell type-specific consequences for cytokinesis failure. Animals from worms to humans can have organism-wide genetic mutations that result in highly cell type-specific cytokinetic consequences.^15,17-35^ On one hand, cell type-specific failure in cytokinesis, resulting in a binucleated tetraploid cell, is emerging as an important contributor to many diseases including blood disorders, neurological diseases, and cancer.^17,23,27,30,33,36-43^ On the other hand, cytokinesis failure is not always pathogenic and specific cell types (*e.g.*, hepatocytes in the liver and intermediate cells in the bladder) are naturally programmed to fail in cytokinesis and become bi-nucleated (or multi-nucleated) as a normal part of human development and tissue homeostasis.^36,44-50^ Despite this strong supporting evidence of cell type-specific regulation of cytokinesis, the molecular mechanisms remain poorly understood.

In theory, the molecular mechanisms differentially regulating cytokinesis in different cell types should arise in the literature as molecules that differentially affect cell division in different cell types and model systems. Two such candidates for cell type-specific regulation of cytokinesis are the septins and anillin. Septins and anillin are cytoskeletal-binding proteins essential for cytokinesis in some, but not all, cell types and model systems; their precise roles in cytokinesis remain unclear.^51-53^ The septins are essential for cytokinesis in budding yeast^54,55^ but are not required for cytokinesis in other cell types, including in *S. pombe^56^*, mouse myeloid and lymphoid hematopoietic cells^25^, and mouse neuronal precursor cells^57^. And, in cultured mammary epithelial cells, *septin-6* expression is inversely correlated with successful cytokinesis, suggesting an inhibitory role.^58^ Similarly, anillin is required for cytokinesis in some cell types^59-61^, including in the fission yeast *S. pombe*^62-64^, *Drosophila* S2 cells^61,65^, and HeLa cells^61^, but not required in many cell types, including the fission yeast *S. japonicus*^66^. In *S. cerevisiae*, anillin (Boi1/2p) is only required in the presence of DNA bridges in the cell division plane.^67,68^ In *C. elegans*, neither the septins nor anillin^ANI-1^ are required for early cleavage divisions during embryogenesis.^69,70^ Even in one-celled organisms that require septins and anillin, these proteins have different functions and localization than in multicellular organisms. For example, the septin ring at the bud neck in *S. cerevisiae* separates into two rings that sandwich, rather than overlap with, the actomyosin contractile ring prior to ring constriction^71,72^, suggesting that septins need to get out of the way for cytokinesis to proceed. Likewise, in *S. pombe*, anillin (Mid1p) leaves the division plane before contractile ring constriction.^73^ These data suggest a cell type- and model system-specific role for septins and anillin in cytokinesis.

We hypothesize that septins and anillin play a key role in cell type-specific differences in cytokinesis, particularly in protection of cytokinesis of the germ lineage from damage to the actin cytoskeleton. In previous work, we found that septin^UNC-59^ and anillin^ANI-1^ likely act as negative regulators of mitotic cytokinesis in 1-cell *C. elegans* embryos.^74^ We also identified cell type-specific regulation of cytokinesis at the 4-to-8-cell stage for the germ precursor cell. We damaged the actin cytoskeleton using either a genetically encoded fast-acting temperature sensitive (ts) mutant that affects the filamentous actin (F-actin) nucleating activity^75^ of the diaphanous family formin^CYK-1^ (hereafter, *formin(ts)*), or a chemical inhibitor of F-actin assembly, Latrunculin A. Under both conditions, the two anterior cells (ABa and ABp) always failed in cytokinesis, whereas the two posterior cells (EMS and P2) divided successfully at a high frequency, even without detectable F-actin in the cell division plane.^15^ Interestingly, we found that cytokinetic protection of EMS and P2 is regulated by a distinct molecular mechanism in each cell. Using embryo micro-dissection to physically separate each of the 4 cells from *formin(ts)* embryos, only the P2 cell germ precursor cell was still protected from cytokinesis failure; EMS lost its protection and failed to divide.^15^ Thus, cell type-specific protection of cytokinesis in the P2 germ precursor cell is cell-intrinsic and in the EMS cell it is cell-extrinsic.

Here, to examine the cell type-specific regulation of cytokinesis that underlies cell-intrinsic protection of germ precursor cells, we examined the role of germ cell fate determinants in cytokinesis. Three well-established and essential germ cell fate determinants are MEX-1, PIE-1, and POS-1, all of which encode CCCH Zn-finger proteins.^76-78^ PIE-1 (pharynx and intestine in excess) is a master regulator of germ cell fate specification in worms.^1,76,79-81^ PIE-1 is asymmetrically inherited by the germ precursor cells where it localizes to ribonucleoprotein condensates called germ granules (or p-granules) and to the nucleus during interphase, but during mitosis it relocalizes to the centrosomes.^76,82,83^ Several other CCCH Zn-finger proteins, including POS-1 and MEX-1, cooperate to control proper PIE-1 localization in germ precursor cells^77,84^ and residual PIE-1 protein (and POS-1 and MEX-1) in somatic daughters is degraded via proteolysis in an E3 ligase substrate adaptor (ZIF-1)-dependent manner^85,86^. PIE-1 canonically controls germ fate specification by regulating transcription^87-91^, gene silencing^92^, translation^84^, and post-translational modifications (*e.g.*, acetylation and SUMOylation)^92^ through inhibition of a NuRD (nucleosome remodeling and deacetylase) complex^92,93^. Neither PIE-1, nor other CCCH Zn-finger proteins that regulate germ fate in *C. elegans*, have previously been implicated in cytokinesis.

We show that protection of P2 cytokinesis is tied to its cellular identity as a germline precursor cell. While another group recently reported a positive role for anillin^ANI-1^ during cytokinesis when central spindle assembly is disrupted in the EMS cell at the 4-cell stage^94^, our research aligns more closely with instances in numerous animal cell types in which septins and anillin are present at the division plane but are not required for cytokinesis. Here we provide evidence that septins and anillin not only act as negative regulators of cytokinesis but also are controlled by germ cell fate determinants that promote cytokinetic protection. The totipotent P2 germ precursor cell is required to produce all gametes (oocytes and sperm) in the adult worm^95^. We identify three germ fate determinants required for protection of P2 cytokinesis in *formin(ts)* embryos. Depletion of either MEX-1, PIE-1, or POS-1 led to loss of cytokinetic protection and P2 cytokinesis failure in *formin(ts)* embryos, but not in control embryos. Depletion of MEX-1 also led to EMS cytokinesis failure, whereas PIE-1 and POS-1 acted in a P2 cell-specific way. We found that PIE-1 does not appear to play a major role in controlling many factors known to affect cytokinesis, including cell surface tension, spindle dynamics, and asymmetric cell division. Instead, our analysis revealed that these germ fate determinants protect cytokinesis by blocking the excessive accumulation of both septin^UNC-59^ and its binding partner, anillin^ANI-1^, at the P2 cell division plane. Co-depletion of septin^UNC-59^ and PIE-1 (or POS-1) was necessary and sufficient to both reduce anillin^ANI-1^ levels at the P2 division plane and restore cytokinetic protection of P2 in *formin(ts)* embryos. Thus, germ fate specification protects the P2 germ precursor cell from cytokinesis failure upon damage to the actin cytoskeleton at least in part by restricting the levels of septin^UNC-59^ and anillin^ANI-1^ at the P2 division plane.

## Results

### Protection of cytokinesis in the P2 cell requires the germ cell fate determinants MEX-1, POS-1, and PIE-1

To identify genes required to protect the P2 germ precursor cell (**Figure 1A**) against cytokinesis failure upon damage to the actin cytoskeleton, we performed a targeted mini-screen of candidate genes either implicated in germ fate regulation in the literature or differentially expressed in the P2 cell by single cell transcriptomics^96^. Embryonic lethality at permissive temperature was used as a proxy for effective gene knockdown (when applicable, see **Figure S1B**). To weaken the actin cytoskeleton, we used the temperature sensitive *formin^cyk-1^(or596ts)* mutant (hereafter *formin(ts)*), which completely blocks cytokinesis in the 1-cell embryo at restrictive temperature with little to no contractile ring constriction or detectable F-actin in the division plane.^15,75^ P2 cytokinesis was monitored by time-lapse spinning disc confocal microscopy in embryos expressing fluorescently tagged reporters for the plasma membrane and chromatin (GFP::PH^pLCδ^ and mCherry::histone H2B^HIS-58^, respectively^97^; **Figure 1B**). Four-cell stage *control* and candidate feeding RNAi-treated *formin(ts)* embryos were upshifted from 16°C (permissive temperature) to ∼24.5-25.5°C (semi-restrictive temperature) prior to anaphase onset in the P2 cell (**Figure 1B-C**). In untreated *formin(ts)* control embryos at this temperature, while ABa and ABp were unable to divide (0% cytokinesis completion), many EMS and P2 cells completed cytokinesis successfully (68% EMS and 50% P2 cytokinesis completion, respectively; **Figure S1A**). The P2 cell in *control* (empty vector) RNAi-treated embryos also frequently completed cytokinesis successfully (78% P2 cytokinesis completion, **Figure 1C**). While RNAi of most candidate genes had no effect on the success rate of P2 cytokinesis, we identified three genes required for P2 cytokinetic protection in *formin(ts)* embryos. Specifically, RNAi-mediated knockdown of MEX-1, PIE-1, and POS-1 significantly decreased the rate of cytokinesis completion in *formin(ts)* P2 cells (15%, 28%, and 27% cytokinesis completion, respectively; **Figure 1C**). These proteins are all CCCH Zn-finger family members essential for proper germ fate specification but not previously implicated in cytokinesis. This result suggests that cytokinetic protection of P2 depends on the germ fate determinants MEX-1, PIE-1, and POS-1.

**Figure 1:**
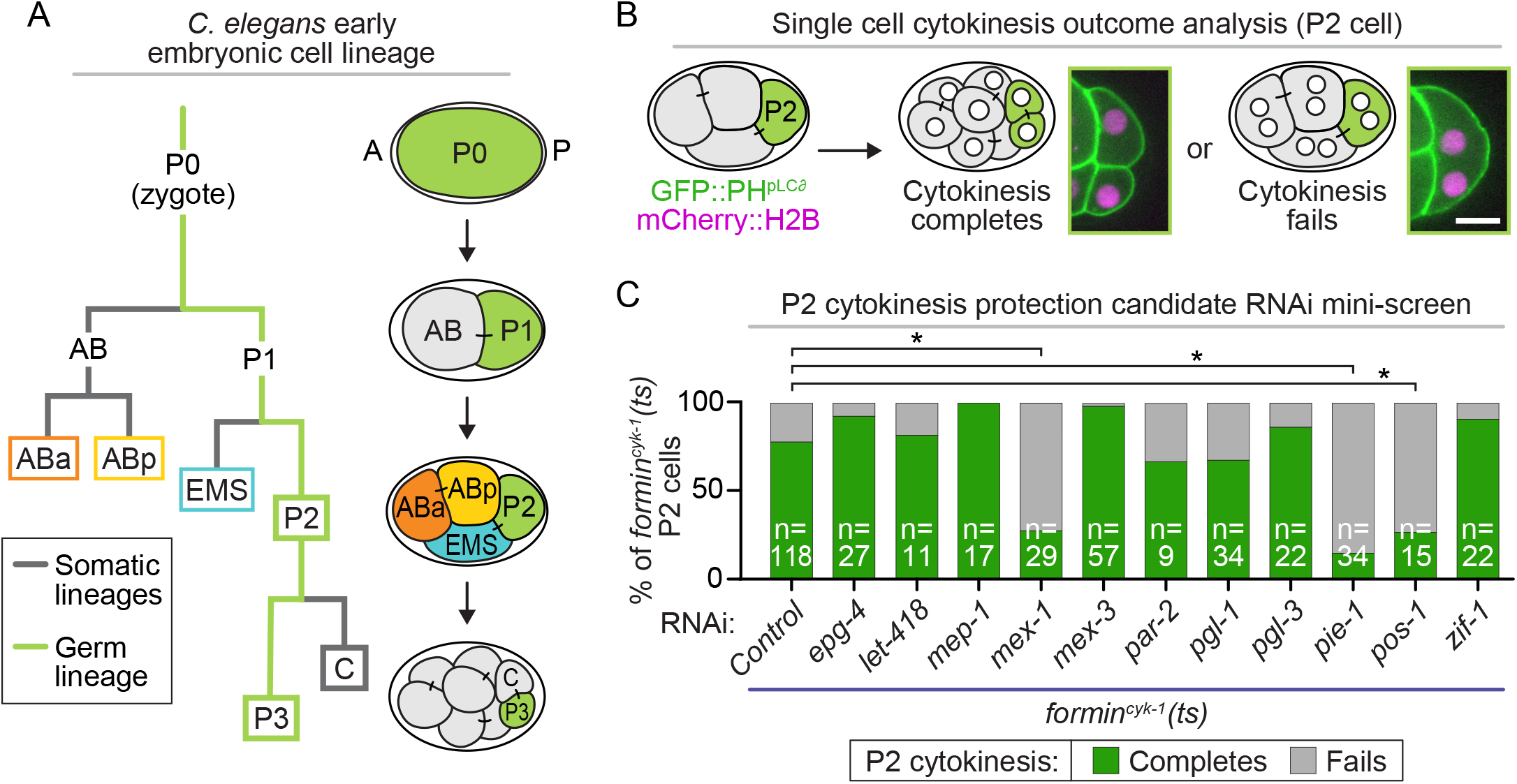
RNAi-based mini-screen of candidate genes required for protection of P2 cytokinesis in formin^cyk-1^(ts) embryos. **A)** Schematic of *C. elegans* early embryonic cell lineage map from the 1- to 4-cell stage (8-cell stage for germ lineage); A=anterior of embryo; P=posterior of embryo; gray lines=somatic cell lineages, lime green lines=germ lineage. **B)** Schematic of single cell cytokinesis outcome analysis for the P2 cell. Representative single plane images show the result of P2 cytokinesis completion (left, 2 mononucleated daughter cells) or failure (right, 1 binucleated daughter cell) in *formin^cyk-1^(ts)* embryos expressing GFP::PH^pLCδ^ (green, plasma membrane) and mCherry::histone H2B^HIS-58^ (magenta, chromatin); scale bar=10 μm. **C)** Graph showing the percentage of P2 cells in *formin^cyk-1^(ts)* embryos that complete (green) or fail (gray) in cytokinesis with or without feeding RNAi treatment; n=number of P2 cells scored and is indicated on each bar; *=p-value ≤0.05 (Fisher’s exact test; see also **Table S1**).

We next sought to determine if this CCCH Zn-finger family-mediated protection of P2 cytokinesis is cell type-specific. We upshifted 4-to-8 cell control and *formin(ts)* embryos prior to anaphase onset in each of the individual 4 cells with and without MEX-1, PIE-1, or POS-1 RNAi-treatment and monitored cytokinesis, as above. For these experiments (and hereafter) we switched to injection RNAi, which is more robust than feeding RNAi in our hands. RNAi knockdown was confirmed by both loss of fluorescent signal in 4-cell embryos expressing fluorescently-tagged reporters of these CCCH Zn-finger proteins (<1%, 4.7%, and <1% of control levels of GFP::MEX-1^98^, GFP::PIE-1^99^, and POS-1::GFP^100^, respectively; **Figure S2A-D**) and consistently high (>99%) embryonic lethality (**Figure S2E-G**). In control embryos, RNAi-mediated knockdown of MEX-1, PIE-1, or POS-1 did not affect cytokinesis in any cell of the 4 cells (100% cytokinesis completion; **Figure 2A**), as predicted^78,79^. In *formin(ts)* embryos, knockdown of MEX- 1, PIE-1, or POS-1 did not change the high rate of cytokinesis failure in the ABa or ABp cells (0% cytokinesis completion in both ABa and ABp; **Figure 2B**) but led to a high frequency of P2 cytokinesis failure (0%-8% cytokinesis completion; **Figure 2B-C**). RNAi-knockdown of MEX-1, but not PIE-1 or POS-1, also led to a high frequency of EMS cytokinesis failure in *formin(ts)* embryos (0% EMS cytokinesis completion in *mex-1(RNAi)*; **Figure 2B**). Together, these results suggest that PIE-1 and POS-1 provide cell type-specific cytokinetic protection of P2, whereas MEX-1 protects cytokinesis in both P2 and EMS.

**Figure 2:**
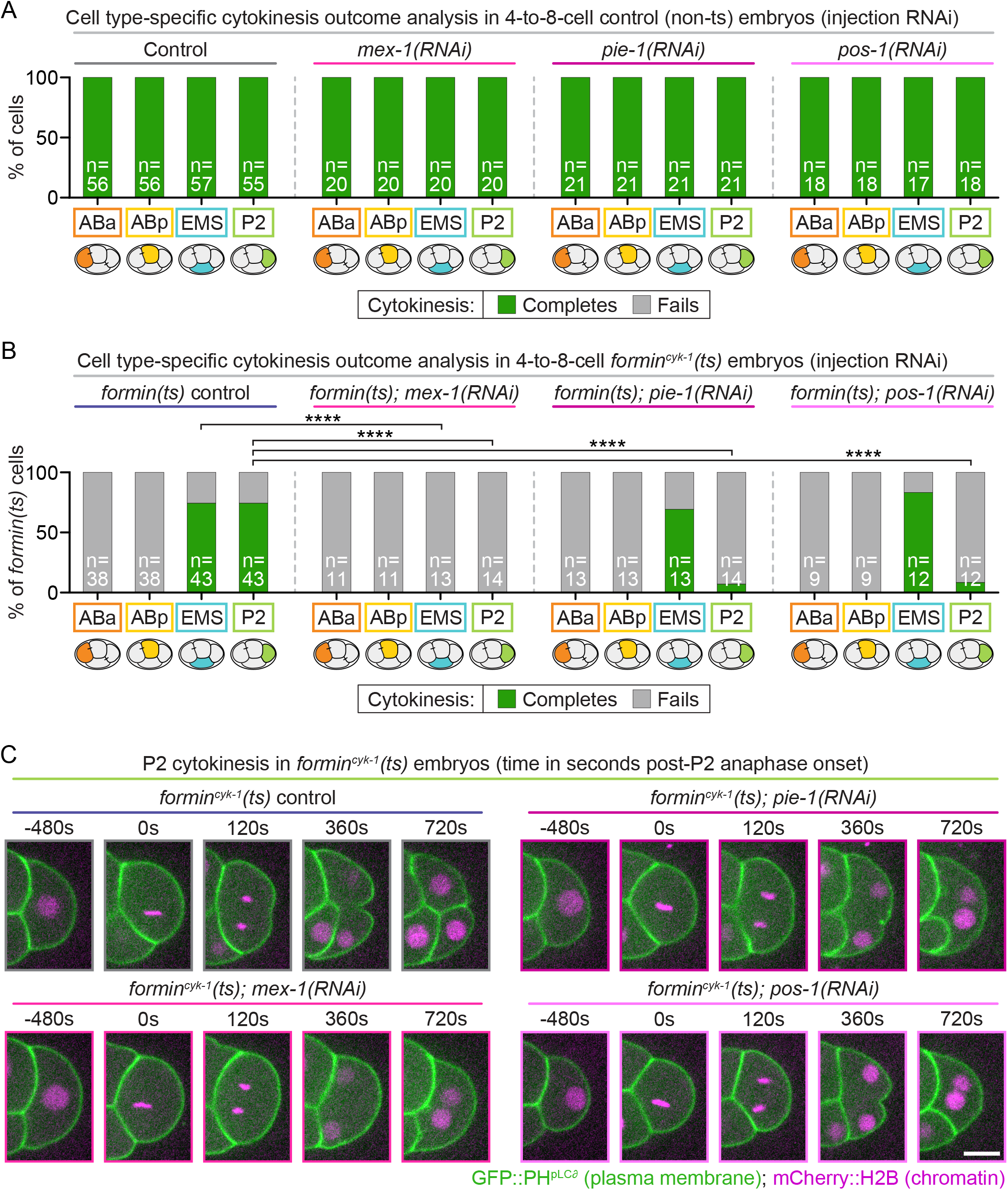
PIE-1 and POS-1 function cell type-specifically to protect cytokinesis in P2, whereas MEX-1 protects cytokinesis in both P2 and EMS. Graphs showing the percentage of ABa, ABp, EMS, and P2 cells that complete (green) or fail (gray) cytokinesis in 4-to-8-cell **A)** control embryos and **B)** *formin^cyk-1^(ts)* embryos with and without *mex-1, pie-1,* or *pos-1* injection RNAi treatment; n=number of individual cells scored and is indicated on each bar; ****=p-value ≤0.0001 (Fisher’s exact test; see also **Table S1**). **C)** Representative single plane images showing P2 cytokinesis in *formin^cyk-1^(ts)* embryos with and without RNAi-mediated depletion of *mex-1, pie-1,* or *pos-1* in embryos expressing GFP::PH^pLCδ^ (green, plasma membrane) and mCherry::histone H2B^HIS-58^ (magenta, chromatin); time (s) is relative to anaphase onset in each P2 cell; scale bar=10 μm.

### PIE-1 does not affect relative surface tensions in P2

It was recently reported that in 4-cell stage embryos, cell surface tension is higher in the anterior ABa and ABp cells than in the posterior EMS and P2 cells.^101^ Because we found that these posterior cells were protected against cytokinesis failure after actin damage and that in P2 cytokinetic protection required key CCCH Zn-finger proteins, we hypothesized that this cytokinetic protection might be mediated by CCCH Zn-finger protein effects on P2 surface tension. We focused on PIE-1 for this analysis because MEX-1 and POS-1 are known to be required for proper PIE-1 localization.^77,78,83,86^ To estimate the relative contribution of surface tension to different success rates of P2 cytokinesis, we measured the contact angles for both the P2-ABp and P2- EMS cell contacts in *formin(ts)* embryos with and without *pie-1(RNAi)* throughout the P2 cell cycle up until the onset of furrowing and used a Young-Dupré force balance to estimate surface tension ratios, as was done previously^101^ (**Figure 3A**, see also Methods). We found no significant difference in the relative tensions of P2 surfaces between control and *pie-1(RNAi)* embryos (**Figure 3B-E**). Cytokinesis failure was associated with altered tension patterns in the embryo (in ABp and/or P2 cells; **Figure S3A**), but the precise tension patterns associated with successful cytokinesis seem to be distinct in control and *pie-1(RNAi)* embryos (**Figure S3B, C**). Together, our results do not suggest a major role for PIE-1 in regulating P2 surface tension.

**Figure 3:**
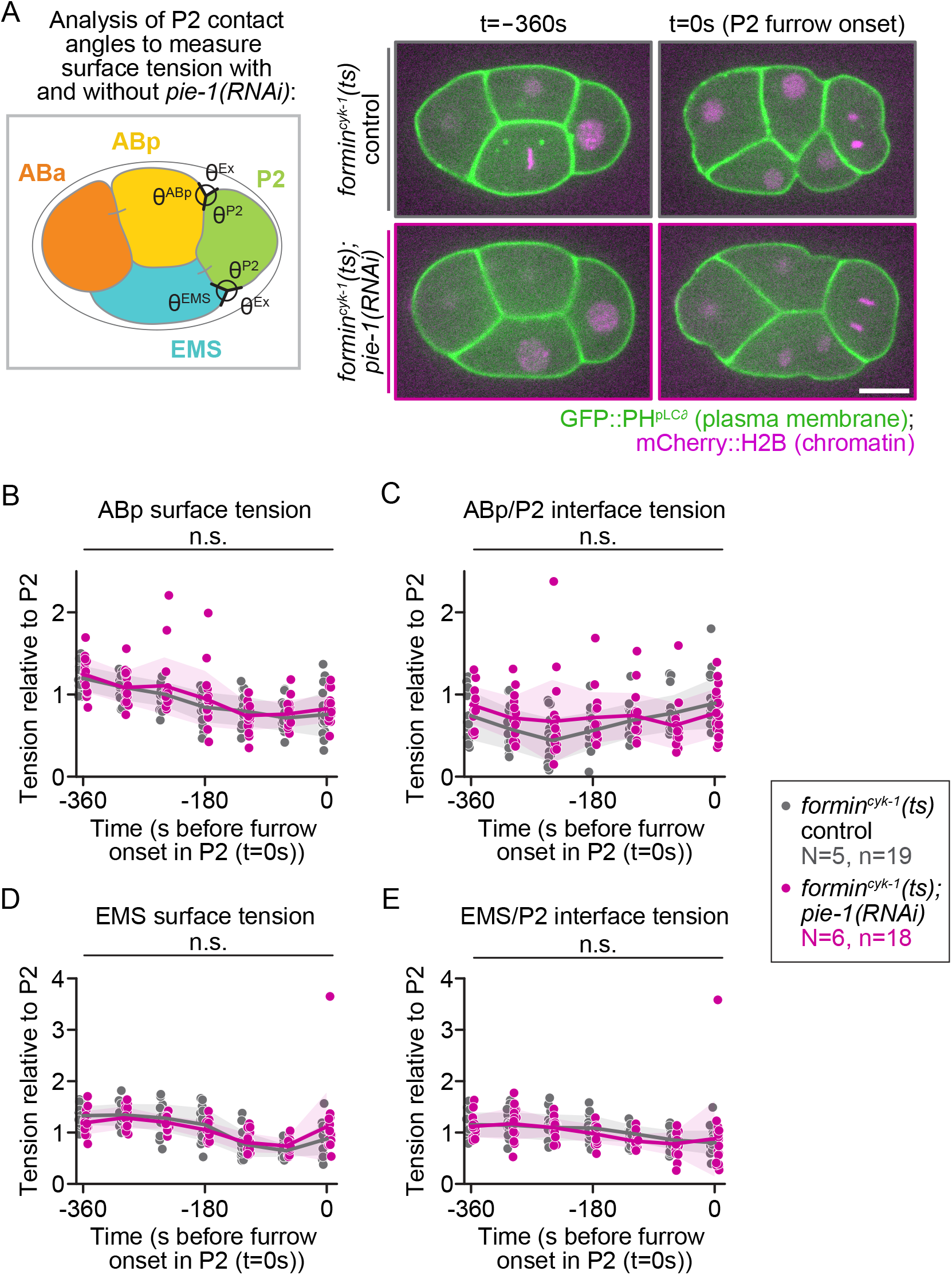
PIE-1 does not significantly change relative P2 cell surface tensions. **A)** Schematic (left) and representative images (right) depicting how cell-cell contact angles with P2 (lime green) neighboring cells ABp (yellow) and EMS (teal) were measured (black angles) over time to calculate the relative cell surface and cell-cell interface tension in *formin^cyk-1^(ts)* control (upper panels, gray) and *formin^cyk-1^(ts); pie-1(RNAi)* embryos (lower panels, pink) expressing GFP::PH^pLCδ^ (green, plasma membrane) and mCherry::histone H2B^HIS-58^ (magenta, chromatin) prior to P2 cytokinesis (similar to as was done in^101^; tension measurements taken on *formin(ts)* control and *formin(ts); pie-1(RNAi)* datasets in Figure 2B and Figure 8; see also Methods); scale bar=10 μm. Graphs showing the relative **B)** ABp surface tension, **C)** ABp/P2 interface tension, **D)** EMS surface tension, and **E)** EMS/P2 interface tension in control (gray) and *pie-1(RNAi)* (pink) embryos. Tension measurements normalized to P2 surface tension; time (s) is relative to furrow onset (t=0s) in each P2 cell; N=number of experimental replicates; n=number of embryos scored for each genotype by color; n.s.=p-value not significant (Student’s t-test, unpaired with Holm-Sidak correction; see also **Table S1**).

### PIE-1 does not affect overall spindle dynamics but has a minor effect on P2 cell and spindle size

Because signals from anaphase spindle microtubules are critical for cytokinesis in animal cells^102,103^, we tested if PIE-1 mediates cytokinetic protection by regulating P2 spindle dynamics. During interphase, PIE-1 localizes to the nucleus and specialized ribonucleoprotein germ granules in the cytoplasm; during mitosis, PIE-1 localizes asymmetrically at P2 spindle poles and is enriched on the germ daughter P3-destined centrosome relative to the somatic daughter C- destined centrosome.^76,83^ To test if PIE-1 regulates the P2 anaphase spindle, we first examined overall spindle and cellular dynamics throughout P2 cell division (**Figure 4A, Figure S4A**) with and without *pie-1(RNAi)* in a strain expressing fluorescently-tagged reporters to label the centrosomes (EB1^EBP-2^::GFP^104^), chromatin^97^, and plasma membrane^9^. We found small differences in overall P2 spindle and cellular dynamics. Relative to dividing P2 cells in controls, P2 cells in *pie-1(RNAi)* embryos had a slight but significantly increased P2 spindle length (centrosome to centrosome distance, **Figure 4B**), P2 cell length, P2 division plane diameter, and diameter of both forming C and P3 daughter cells (**Figure S4B-E**). The distance from the anterior cell cortex to the germ daughter P3-destined centrosome was also significantly increased in *pie-1(RNAi)* relative to control embryos, but there was no difference in the distance from the posterior cell cortex to the somatic daughter C-destined centrosome (**Figure S4F, G**). There were only minor differences in the separation of sister chromosomes in anaphase in *pie-1(RNAi)* relative to control embryos (**Figure 4C**). We next tested if PIE-1 regulates P2 astral microtubule dynamics by imaging the EB1^EBP-2^::GFP microtubule plus-tip binding protein at higher temporal resolution. We found no difference in astral microtubule growth rates in P2 anaphase from either the C- or P3-destined centrosomal asters with and without *pie-1(RNAi)* (**Figure 4D**). Finally, we assessed central spindle assembly in control and *pie-1(RNAi)* embryos. The central spindle is an antiparallel microtubule structure that forms between separating chromatids in anaphase and also plays a role in contractile ring constriction.^102,103^ Using a reporter for central spindle assembly (Aurora-B^AIR-2^::GFP^105^)^106,107^, we found no difference in the timing or morphology of central spindle assembly in *pie-1(RNAi)* versus in control P2 cells (**Figure 4E-F, Figure S5**). Thus, PIE-1 has minor effects on P2 cell and spindle size, but despite localizing to the centrosomes, does not seem to have any major effects on overall P2 anaphase spindle dynamics.

**Figure 4:**
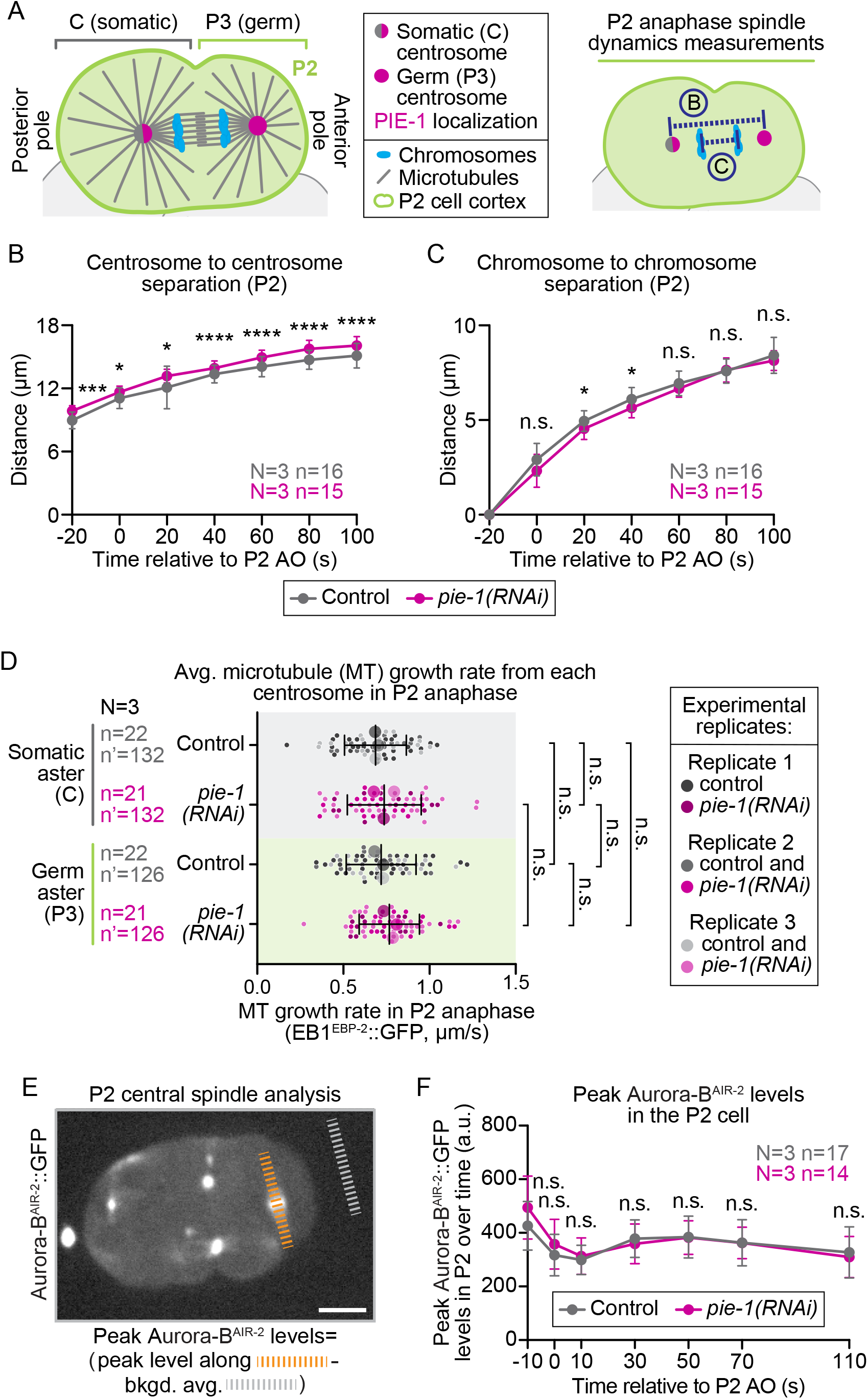
PIE-1 does not significantly alter P2 spindle or daughter cell dynamics. **A)** (Left) Schematic of a dividing P2 cell indicating the cell cortex (lime green), chromosomes (blue), centrosomes (C centrosome (half gray/half pink); P3 centrosome (pink)), and microtubules (gray). (Right) Schematic of measurements taken and plotted in **B** and **C** (see also Methods). Graphs plotting the kinetics of **B)** centrosome separation and **C)** chromosome separation over time (s) relative to anaphase onset (AO, t=0s) in dividing control (gray) and *pie-1(RNAi)* (pink) P2 cells expressing EB1^EBP-2^::GFP, mCherry::histone H2B^HIS-58^, mCherry::PH^pLCδ^. Error bars=standard deviation; N=number of experimental replicates; n=number of P2 cells scored for each genotype by color; n.s.=not significant, *=p-value ≤0.05, ***=p-value ≤0.001, and ****=p- value ≤0.0001 (Student’s t-test, unpaired, see also **Table S1**). **D)** Graph showing super plots of the average growth rates for EB1^EBP-2^::GFP-labeled astral microtubules (MTs) emanating from the somatic-(C) or germ-daughter cell (P3) destined centrosome in dividing control (grays) and *pie-1(RNAi)* (pinks) P2 cells at 26°C. Small circles indicate individual data points and large circles and color shades indicate replicate averages; error bars=standard deviation; N=number of experimental replicates; n=number of P2 cells scored; n’=number of astral microtubules scored for each genotype by color; n.s.=not significant (two-way ANOVA, see also **Table S1**). **E)** Representative images (maximum projection) of a 4-to-8-cell embryo expressing Aurora-B^AIR-2^::GFP and mCherry::H2B (not shown, see **Figure S5**) depicting linescan analysis used on sum projected embryos to quantify central spindle (orange dashed line) and camera background (gray dashed line) shown in **F**; scale bar=10 μm. **F)** Graph plotting the average peak Aurora-B^AIR-2^::GFP levels at chromosomes (pre-anaphase onset, AO (metaphase)) and the central spindle (post-AO) during P2 cell division in control (gray) and *pie-1(RNAi)* (pink) P2 cells over time. Time (s) is relative to anaphase onset (AO, t=0s) in each P2 cell; error bars=standard deviation; N=number of experimental replicates; n=number of P2 cells scored for each genotype by color; n.s.=not significant (Student’s t-test, unpaired, see also **Table S1**).

### PIE-1 plays a minor role in regulating daughter cell asymmetry during P2 cell division

Given the small differences in overall P2 and daughter cell size we observed during cell division with and without PIE-1, we tested if PIE-1 affects P2 division asymmetry. The P2 cell divides asymmetrically, producing a larger somatic precursor daughter cell in the posterior and a smaller germ precursor daughter cell in the anterior.^4,108,109^ PIE-1 is asymmetrically inherited by the germ precursor cells throughout early worm development^4,76,82,110^, but is not thought to regulate cell division asymmetry directly. Myosin-II^NMY-2^ is a key regulator of asymmetric cell division in *C. elegans*.^108,111-114^ To test if PIE-1 affects P2 cytokinesis by regulating polarity, we monitored polarity by measuring peak levels of myosin-II^NMY-2^::GFP on the cortex in both forming daughter cells during P2 cytokinesis with and without *pie-1(RNAi)* (**Figure 5A-B**). In control embryos, we found higher levels of myosin-II^NMY-2^ on the C-destined daughter cell cortex than on the P3-destined daughter cell cortex (**Figure 5C**), as would be predicted^108,109,112^. In *pie-1(RNAi)* embryos, the levels of myosin-II^NMY-2^ were also higher on the C-destined daughter cell cortex than on the P3-destined daughter cell cortex, although to a lesser extent than in control embryos (**Figure 5C**). These results suggest that PIE-1 plays a minor role regulating myosin-II^NMY-2^ on the germ daughter-destined cell cortex but is not essential for overall cortical asymmetry during P2 cell division.

**Figure 5:**
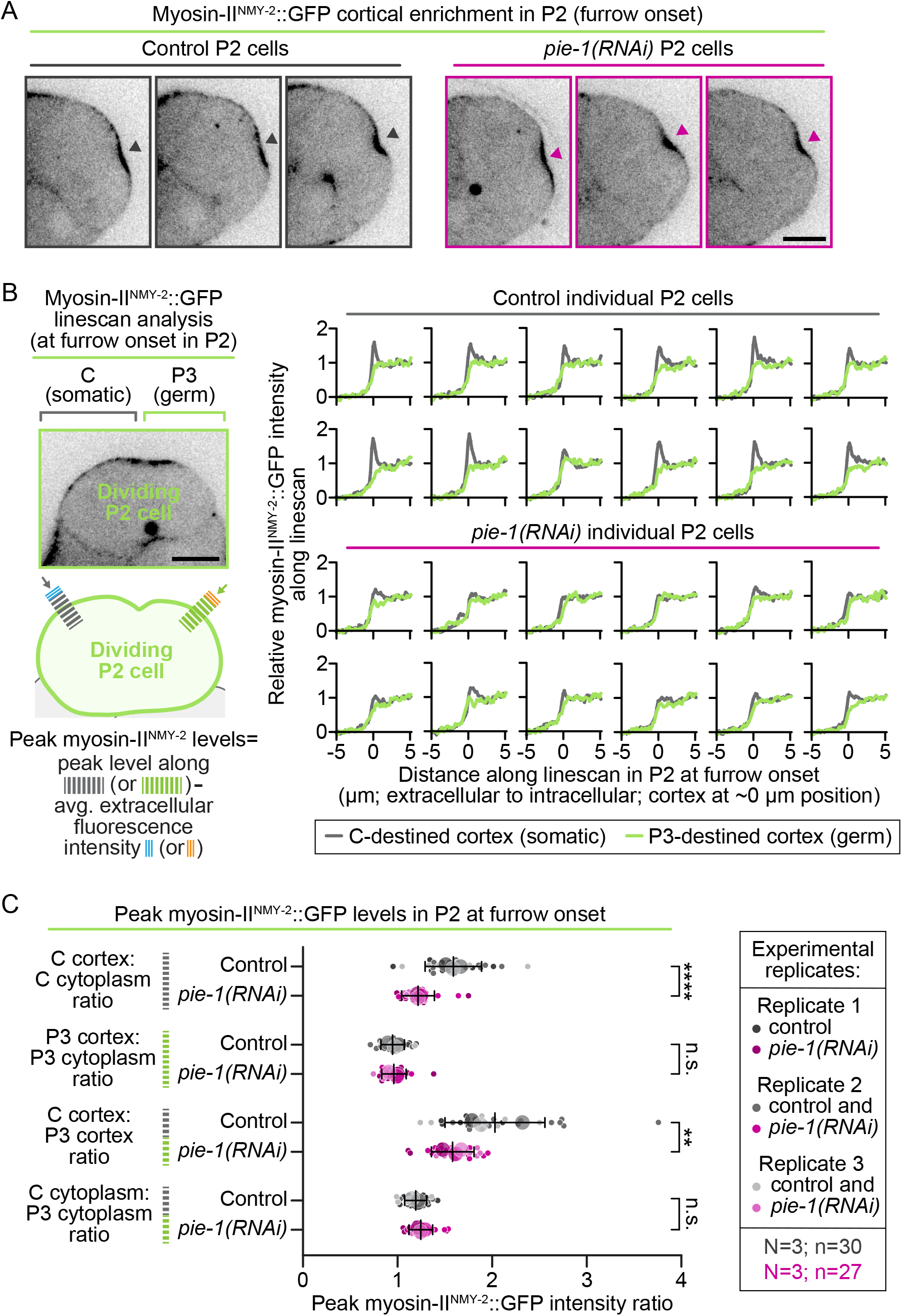
PIE-1 plays a minor role in P2 division asymmetry. **A)** Representative single plane images of 3 control (left) and 3 *pie-1(RNAi)* (right) P2 cells expressing endogenously-tagged myosin-II^NMY-2^::GFP at the time of cleavage furrow onset in P2; scale bar=10 μm; gray (control) and pink (*pie-1(RNAi)*) arrowheads indicate the P2 cleavage furrow. **B)** (Left) Schematic depicting linescan analysis used to quantify cortical asymmetry of myosin-II^NMY-2^ in the P2 cell during division (image, top); arrows indicate direction of linescans drawn across the C-destined and P3-destined cell cortices from the respective extracellular to intracellular space (schematic, bottom; see also Methods). (Right) Representative linescans from 12 control (top) and 12 *pie-1(RNAi)* (bottom) P2 cells plotting relative myosin-II^NMY-2^::GFP levels along linescans across the C-destined (gray) and P3-destined (lime green) cell cortices. **C)** Graph plotting peak cortical intensity ratios for myosin-II^NMY-2^::GFP in control (gray) and *pie-1(RNAi)* (pink) embryos at the forming C and P3 daughter cell cortexes (relative to average cortical and cytoplasmic levels) during cell division. Error bars=standard deviation; N=number of experimental replicates; n=number of embryos scored for each genotype by color; n.s.=not significant, **=p- value ≤0.01, and ****=p-value ≤0.0001 (Student’s t-test, unpaired, see also **Table S1**).

In the 1-cell embryo, we previously found that the cell polarity machinery was required to sequester anillin^ANI-1^ and septin^UNC-59^ on the anterior side of the cell cortex during cytokinesis.^74^ Thus, we also tested if PIE-1 regulates the cortical asymmetry of anillin^ANI-1^ and/or septin^UNC-59^ in the P2 cell. In control embryos, we found higher levels of anillin^ANI-1^ on the posterior C-destined daughter cell cortex than on the anterior P3-destined daughter cell cortex (**Figure S6**), similar to myosin^NMY-2^. In *pie-1(RNAi)* embryos, similar to myosin^NMY-2^, anillin^ANI-1^ levels were also higher on the C-destined daughter cell cortex than on the P3-destined daughter cell cortex, but to a lesser extent than in control embryos (**Figure S6**). We found no detectable enrichment of septin^UNC-59^ on either the C or P3 sides of the dividing P2 cell cortex, with or without PIE-1 depletion (**Figure S6**). This result suggests that cortical septin^UNC-59^ is not asymmetrically distributed during P2 division and, as with myosin-II^NMY-2^ and P2 cell size, PIE-1 may play a minor role in regulating asymmetric anillin^ANI-1^ levels on the somatic daughter-destined cell cortex.

### PIE-1 and POS-1 restrict accumulation of septin^UNC-59^ and anillin^ANI-1^ at the P2 cell division plane

While measuring the cortical asymmetry of contractile ring proteins during P2 cell division, we observed apparent changes in their protein levels at the P2 division plane following PIE-1 depletion. To test this, we quantified the effect of *pie-1(RNAi)* on contractile ring protein levels at the P2 cell division plane. We imaged the P2 contractile ring when the cleavage furrow was first visible by DIC microscopy (furrow onset or 20s post-furrow onset for septin^UNC-59^::GFP) in strains expressing fluorescently-tagged reporters for multiple contractile ring proteins (LifeAct::RFP and plastin^PLST-1^::GFP (F-actin)^115^, myosin-II^NMY-2^::GFP^116^, septin^UNC-59^::GFP^117^, and anillin^ANI- 1^::GFP^118^). Quantitative analysis revealed no significant difference in the levels of F-actin or the motor myosin-II^NMY-2^ at the P2 division plane in control versus in *pie-1(RNAi)* embryos (**Figure 6A-C, F**). In contrast, RNAi knockdown of PIE-1 significantly increased the cortical levels of both septin^UNC-59^ and anillin^ANI-1^ at the P2 division plane relative to those of control embryos (∼22% higher for septin^UNC-59^ and ∼14% higher for anillin^ANI-1^ in *pie-1(RNAi)* embryos than in controls; **Figure 6D-F**). Knockdown of PIE-1 also increased the total 4-to-8-cell whole embryo levels of septin^UNC-59^ (∼16% higher), but not anillin^ANI-1^, F-actin, or myosin-II^NMY-2^ (**Figure S7**). Together, these results suggest that PIE-1, a critical regulator of germ precursor cell fate, also functions to control the contractile ring levels of septin^UNC-59^ and its binding partner anillin^ANI-1^ during P2 cytokinesis.

**Figure 6:**
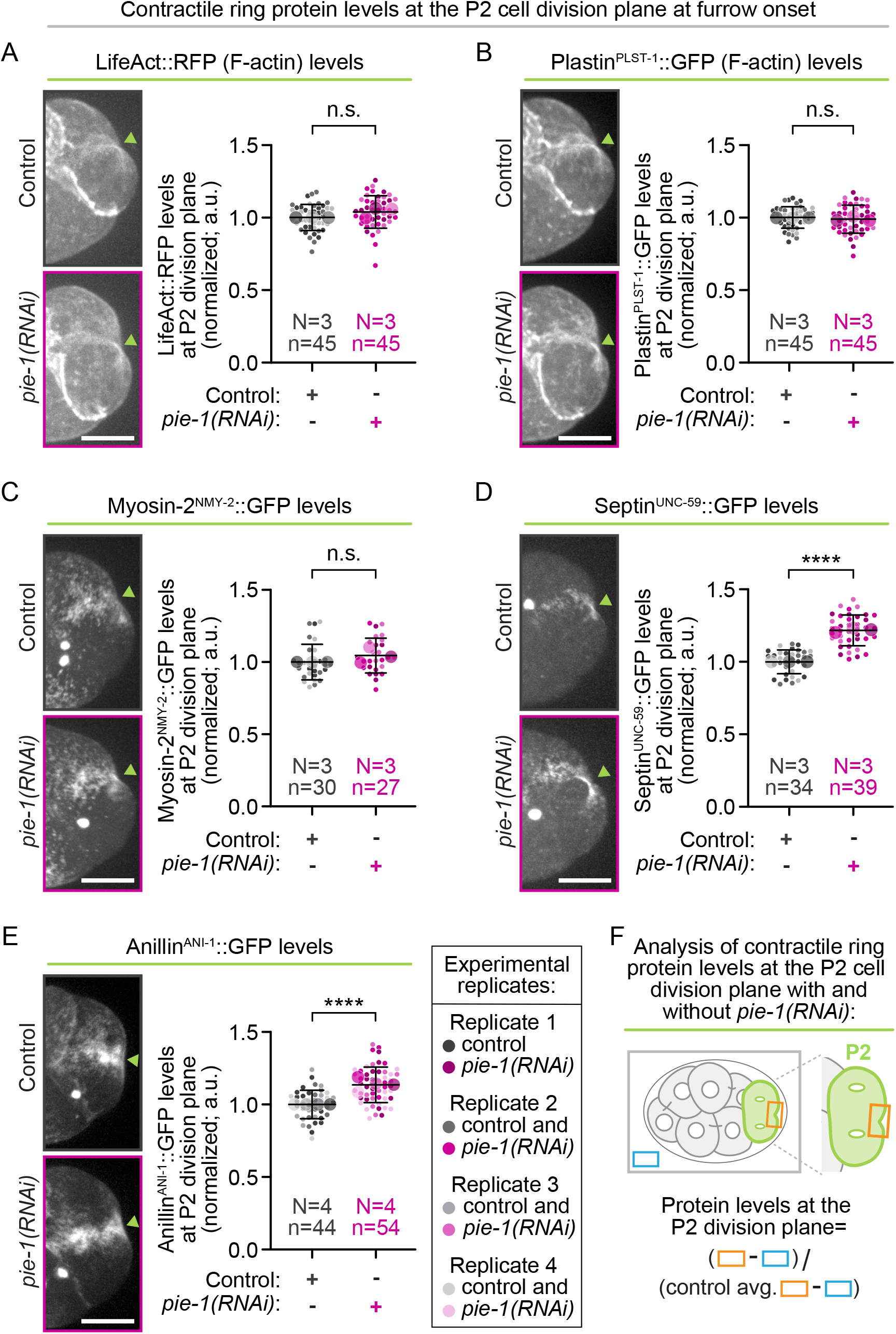
PIE-1 restricts the levels of septin^UNC-59^ and anillin^ANI-1^ at the P2 contractile ring. Representative maximum projection images (left) and graphs (right) showing super plots of normalized P2 contractile ring levels in control (grays) and *pie-1(RNAi)* (pinks) embryos expressing fluorescently tagged reporters for **A, B)** F-actin (**A)** Lifeact and **B)** Plastin^PLST-1^), **C)** myosin-II^NMY-2^, **D)** septin^UNC-59^, and **E)** anillin^ANI-1^ at the time of cleavage furrow onset in P2. Small circles indicate individual data points and large circles and color shades indicate replicate averages; scale bar=10 μm; lime green arrowheads indicate the P2 cleavage furrow; error bars=standard deviation; N=number of experimental replicates; n=number of embryos scored for each genotype by color; n.s.=not significant, ****=p-value ≤0.0001 (Student’s t-test, unpaired, see also **Table S1**). **F)** Schematic depicting analysis shown in **A-E** performed on sum projected images to measure contractile ring protein levels at the P2 contractile ring (orange box) and extracellular background (blue box, see also Methods).

While anillin often functions upstream of septins^13,119^, anillin levels during cytokinesis are regulated by septin in some cellular contexts (*e.g.*, see^14,53,65^). To test whether PIE-1 and/or POS- 1 act through septin^UNC-59^ to regulate the cortical levels of anillin^ANI-1^ at the P2 division plane after actin damage in *formin(ts)* embryos, we imaged the levels of endogenously-tagged^118^ anillin^ANI-1^ in *formin(ts)* embryos with and without RNAi knockdown of PIE-1, POS-1, and septin^UNC-59^ individually and together. RNAi knockdown was confirmed by loss of fluorescent signal in 4-cell embryos expressing fluorescently-tagged reporters of PIE-1^99^, POS-1^100^, and septin^UNC-59^ ^117^ (<1% of control levels in respective RNAi-mediated double knockdown embryos; **Figure S8**). Anillin^ANI-1^ levels at the P2 division plane and in whole 4-to-8-cell embryos were much higher in *formin(ts)* mutants relative to those of control embryos with no ts mutations (**Figure 7, Figure S9**), as was recently reported in formin^CYK-1^-disrupted 1-cell embryos^120^. RNAi knockdown of septin^UNC-59^ reduced anillin^ANI-1^ levels at the P2 division plane, but not total embryo levels, in both control and *formin(ts)* embryos (∼12% lower in control and ∼21% lower in *formin(ts)* embryos; **Figure 7, Figure S9**). RNAi knockdown of PIE-1 increased anillin^ANI-1^ levels at the P2 division plane to a similar extent in control (**Figure 6E**) and *formin(ts)* embryos (**Figure 7**) (∼14% higher levels in control and ∼13% in *formin(ts)* embryos). POS-1 knockdown also led to increased levels of anillin^ANI-1^ at the P2 division plane in *formin(ts)* embryos (∼18% higher levels in *formin(ts); pos-1(RNAi)* embryos than in *formin(ts)* control embryos; **Figure 7**). The increase in anillin^ANI-1^ at the P2 division plane after PIE-1 or POS-1 knockdown was dependent on septin^UNC-59^, as co-depletion of either PIE-1 or POS-1 with septin^UNC-59^ in *formin(ts)* embryos reduced cortical anillin^ANI-1^ levels at the P2 division plane to a similar extent as depletion of septin^UNC-59^ on its own (∼22% lower in *formin(ts); septin^unc-59^(RNAi)*, ∼19% lower in *formin(ts); pie-1(RNAi); septin^unc-59^(RNAi),* and ∼20% lower in *formin(ts)*; *pos-1(RNAi); septin^unc-59^(RNAi)* relative to in *formin(ts)* controls; **Figure 7**). Thus, PIE-1 and POS-1 require septin^UNC-59^ to control the levels of anillin^ANI-1^ in the P2 division plane in *formin(ts)* embryos.

**Figure 7:**
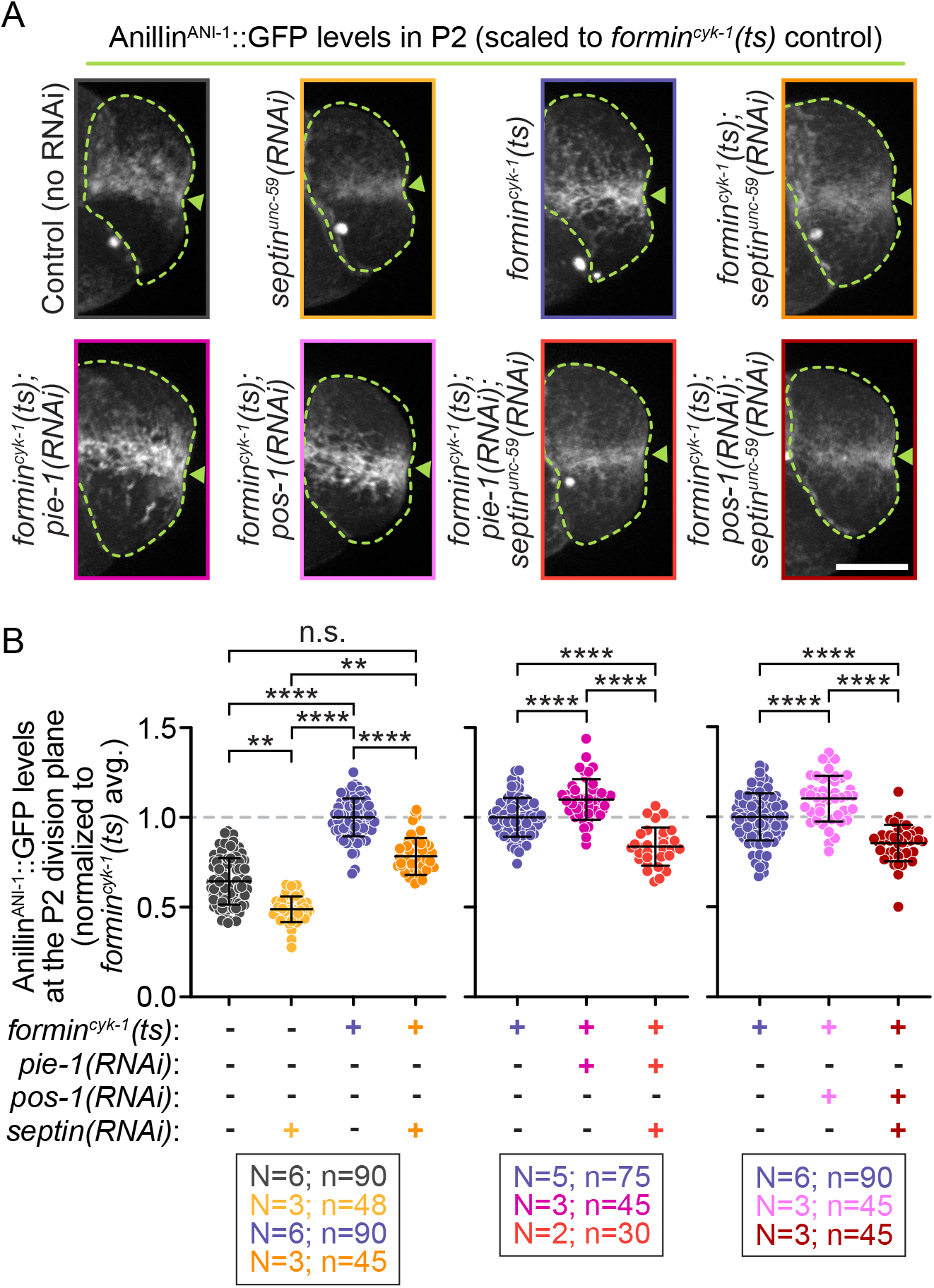
Septin^UNC-59^ is required for PIE-1 to restrict anillin^ANI-1^ levels at the P2 contractile ring. **A)** Representative maximum projection images of endogenously-tagged anillin^ANI-1^::GFP in P2 cells at cleavage furrow onset for indicated genotypes (images scaled relative to *formin^cyk-1^(ts)* control embryos); scale bar=10 μm; lime green arrowheads indicate the P2 cleavage furrow. **B)** Graphs showing normalized anillin^ANI-1^::GFP levels at the P2 division plane for each genotype (normalized to *formin^cyk-1^(ts)* control embryos). Septin^UNC-59^ is required for the increased levels of anillin^ANI-1^ at the P2 division plane in *pie-1(RNAi); formin^cyk-1^(ts)* and *pos-1(RNAi); formin^cyk-1^(ts)* embryos. Error bars=standard deviation; N=number of experimental replicates; n=number of embryos scored for each genotype by color; n.s.=not significant, **=p-value ≤0.01, ****=p-value ≤0.0001 (two-way ANOVA, see also **Table S1**).

### Septin^UNC-59^ depletion is sufficient to restore protection of P2 cytokinesis, even when co-depleted with PIE-1 or POS-1

Given our results, we hypothesized that P2 cytokinetic protection from actin insult is mediated by PIE-1 preventing excess septin^UNC-59^ (and therefore anillin^ANI-1^) accumulation at the division plane. To directly test if reducing septin^UNC-59^ levels could restore P2 cytokinetic protection in *formin(ts)* embryos after loss of PIE-1, we monitored P2 cytokinesis by time-lapse spinning disc confocal microscopy in embryos with and without single and double RNAi treatment, as above (see also **Figure S8**). In *formin(ts)* control embryos (with intact PIE-1 and POS-1), P2 was protected against cytokinesis failure and completed cytokinesis frequently (89% cytokinesis completion; **Figure 8A, B**). P2 also completed cytokinesis frequently in *formin(ts)* embryos after RNAi knockdown of septin^UNC-59^ or anillin^ANI-1^ (92% and 90% cytokinesis completion, respectively; **Figure 8A, B**). Again, this P2 cytokinetic protection in *formin(ts)* embryos was lost after knockdown of PIE-1 or POS-1 and cytokinesis failed at a significantly higher frequency (23% and 31% cytokinesis completion, respectively; **Figure 8A, B**). In contrast, co-depletion of septin^UNC-59^ with PIE-1 or POS-1 in *formin(ts)* embryos was sufficient to rescue the frequency of successful P2 division (92% and 93% cytokinesis completion, respectively) (**Figure 8A, B**), and allow P2 cytokinesis to occur in the absence of detectable F-actin at the division plane (**Figure S10**). Together, our results support a model in which the PIE-1 and POS-1 germ fate determinants mediate P2 cytokinetic protection by reducing the levels of two cytoskeletal proteins that seem to act as negative regulators of contractile ring constriction at the division plane, septin^UNC-59^ and anillin^ANI-1^. This germ fate-driven protection of the P2 germ precursor cell allows cytokinesis to complete successfully, even after damage to the actin cytoskeleton (**Figure 8C**).

**Figure 8:**
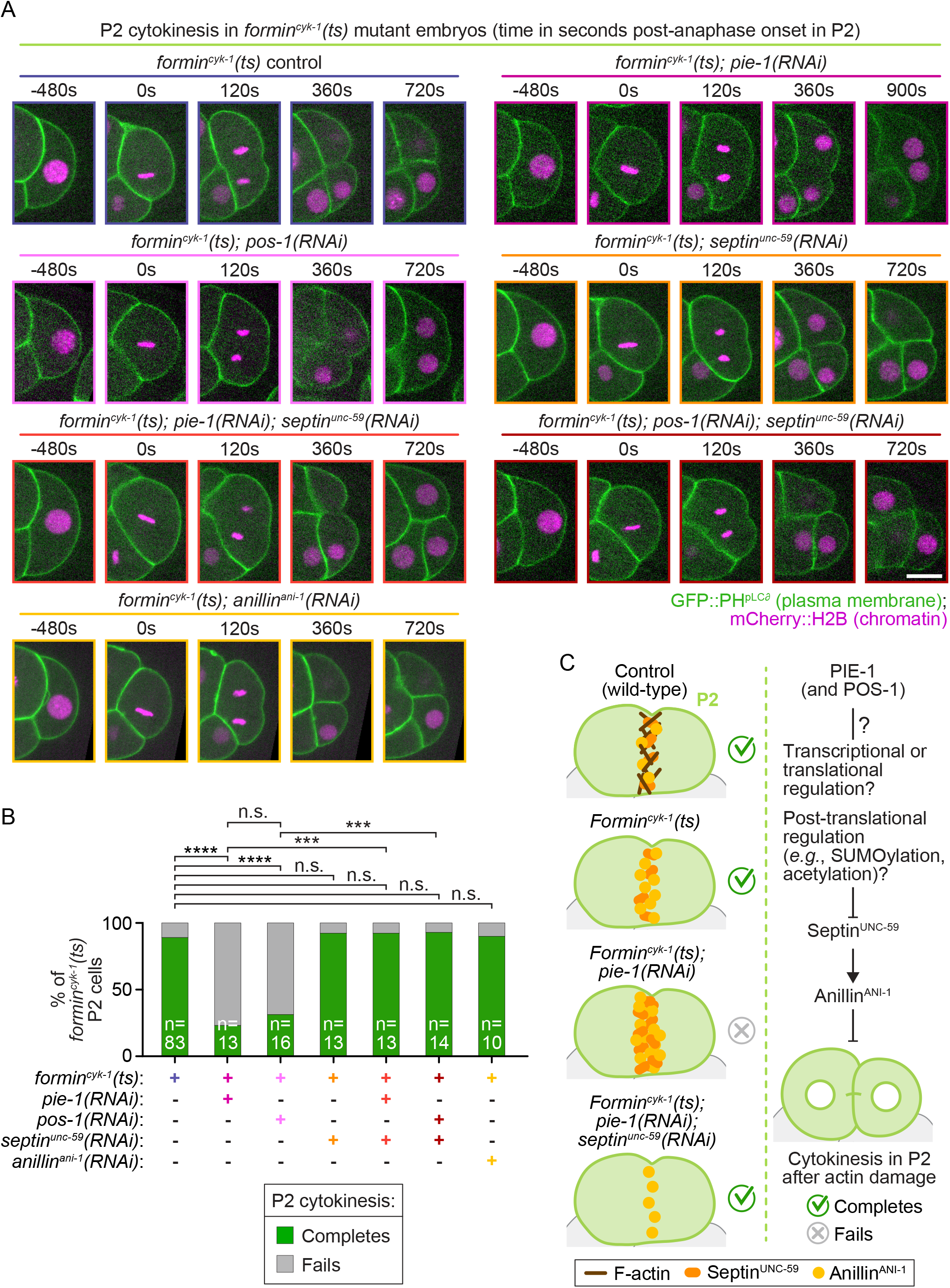
Reducing septin^UNC-59^ levels is sufficient to restore protection of P2 cytokinesis. **A)** Representative single plane images of cytokinesis after indicated RNAi treatment in *formin^cyk-1^(ts)* P2 cells expressing GFP::PH^pLCδ^ (green, plasma membrane) and mCherry::histone H2B^HIS-58^ (magenta, chromatin). Time (s) is relative to anaphase onset in each P2 cell; scale bar=10 μm**. B)** Graphs plotting the percentage of P2 cells in *formin^cyk-1^(ts)* embryos that complete (green) or fail (gray) in cytokinesis with or without indicated injection RNAi treatment. Co-depletion of septin^UNC-59^ restores robust P2 cytokinesis in *formin^cyk-1^(ts); pie-1(RNAi)* embryos. n=number of P2 cells scored and is indicated on each bar; n.s.=not significant, ***=p-value ≤0.001, and ****=p- value ≤0.0001 (Fisher’s exact test, see also **Table S1**). **C)** Schematic (left) and genetic (right) models for the function of PIE-1 and POS-1 in restricting septin^UNC-59^ and anillin^ANI-1^ levels at the P2 division plane to protect cytokinesis after damage to the actomyosin contractile ring.

## Discussion

Here, we investigated the mechanisms that drive cell type-specific protection of cytokinesis in the *C. elegans* P2 germ precursor cell. We identified three well-known germ fate determinants, MEX-1, PIE-1 and POS-1, as required for protection of P2 cytokinesis in *formin(ts)* embryos. Of these, PIE-1 and POS-1 specifically protected cytokinesis in P2, whereas MEX-1 also protected cytokinesis in EMS after actin damage. Neither of these proteins has been previously implicated in cytokinesis and they are not required for cytokinesis in control embryos (*e.g.*, see **Figure 2A**). We found that PIE-1 and POS-1 protect cytokinesis in P2 by preventing excessive accumulation of both septin^UNC-59^ and anillin^ANI-1^, but not F-actin or myosin-II^NMY-2^, at the cell division plane. Co-depletion of septin^UNC-59^ and PIE-1 was both necessary and sufficient to reduce anillin^ANI-1^ levels at the P2 division plane and restore the cytokinetic protection of P2. These data also demonstrate that septin^UNC-59^ and anillin^ANI-1^ can function as negative regulators of cytokinesis. Thus, we found these germ fate determinants protect cytokinesis by controlling the levels of specific actomyosin contractile ring-associated proteins to ensure cytokinesis completes successfully in this germ precursor cell, even without detectable F-actin in the contractile ring.

How do PIE-1 and POS-1 control the levels of septin^UNC-59^ and anillin^ANI-1^ in the P2 germ precursor cell? Canonically, both PIE-1 and POS-1 regulate germ cell fate by controlling transcription and translation.^78,84,86,87,89,90,93,121-124^ The early *C. elegans* germ precursor cells are thought to be transcriptionally silent and PIE-1 is required to maintain transcriptional repression in the germ precursor cells.^83,87,88,90,125^ POS-1 is required for PIE-1 to localize to the nucleus^84^, and thus also likely regulates its transcriptional control. Both POS-1 and PIE-1 are also implicated in translational regulation, and POS-1 is known to regulate poly-A tail length.^122-124^ It is thus possible that germ fate normally leads to transcriptional or translational repression of a negative regulator of cytokinesis (*e.g.*, *septin^unc-59^* and/or *anillin^ani-1^*). Consistent with this idea, the *septin^unc-59^, septin^unc-61^, and anillin^ani-1^* genes all appear to have multiple cytoplasmic polyadenylation elements in their 3”-UTR sequences, suggesting they might be regulated at the translational level. Indeed, in the absence of PIE-1, we observed higher protein levels of endogenously-tagged septin^UNC-59^::GFP both at the P2 division plane and in the whole embryo level. Thus, transcriptional or translational regulation of cytokinesis genes could be controlled by these CCCH Zn-finger proteins.

PIE-1 also regulates post-translational modifications in the germ line (*e.g*., acetylation, SUMOylation^92^) by inhibiting the histone deacetylase HDA-1 and its associate NuRD complex.^92,93,126^ A recent proteomics study revealed that both anillin^ANI-1^ and septin^UNC-59^ are SUMOylated in the worm germline, and disruption of PIE-1 reduces the levels of SUMOylation on both anillin^ANI-1^ and septin^UNC-59^.^92^ Human anillin has also been found to be SUMOylated in a proteomics screen^127^, although the function of anillin SUMOylation is not clear. Septins are SUMOylated in other systems, including in budding yeast, where SUMOylation of several septin proteins (Cdc3, Cdc11, and Shs1) is both cell cycle regulated and required for cytokinesis in this system.^128^ Furthermore, septins from all four human septin groups are also SUMOylated and expression of non-SUMOylatable septin (SEPT6 and SEPT7) variants leads to excessive septin bundling and an increase in multinucleated cells due to a late defect in cytokinesis.^129^ It will be interesting to test if PIE-1-dependent SUMOylation of worm septins^UNC-59/-61^ and/or anillin^ANI-1^ is responsible for protection of P2 cytokinesis.

How does the P2 cell divide without detectable F-actin in the division plane? A recent report suggested that, in the absence of formin^CYK-1^ activity, oligomerized anillin^ANI-1^ itself may interact with myosin-II^NMY-2^ motor proteins and drive cytokinesis in the 1-cell *C. elegans* embryo^120^. However, our finding that the P2 cell in *formin(ts)* mutants could still divide at a high frequency after RNAi knockdown of anillin^ANI-1^ (**Figure** 8A, B) is inconsistent with that model. Moreover, in previous results, we found that inhibition of anillin^ANI-1^ could rescue cytokinesis failure in 1-cell embryos (grandmother cell of P2) after co-disruption of cell polarity and formin^CYK-1^ activity^74^, suggesting a similar mechanism may be at play. These data directly contradict a model in which anillin^ANI-1^ positively drives cytokinesis and are consistent with our proposed model in which septin^UNC-59^ and anillin^ANI-1^ function as negative regulators of cytokinesis, at least in *C. elegans* germ precursor cells.

The fundamental question remains, what is the positive driver of P2 cytokinesis in the absence of F-actin, septin^UNC-59^, and anillin^ANI-1^? We cannot rule out that an adapted filamentous actin system forms upon formin^CYK-1^ disruption that does not associate with LifeAct, plastin^PLST-1^, or utrophin-based^130^ F-actin reporters (**Figure S10**, see also^15^). However, LifeAct binds to highly dynamic F-actin and the utrophin-based reporter binds to more stable actin filaments,^130,131^ so this would have to be a completely non-canonical type of actin filament. To us, it seems more likely that cytokinesis requires significantly lower levels of F-actin (or is F-actin independent) in certain cell type-specific contexts, such as in the P2 cell. Further work will be needed to determine how cells like the P2 germ precursor cell can divide without detectable F-actin in contractile ring.

Cell type-specific protection of cytokinesis after damage to the actin cytoskeleton in cells with a high potency, or ability to differentiate into other cell types, is not necessarily germ precursor cell-specific. Indeed, cell type-specific protection of cytokinesis after actin damage may also apply to human embryonic stem cells. Inhibition of actomyosin contractility is routinely used in *in vitro* cell culture protocols to improve the survival of pluripotent embryonic stem cells, suggesting they can also divide with a weakened contractile ring.^132,133^ Thus, our findings in *C. elegans* germ precursor cells may provide insight into mechanisms of cytokinetic protection in other cell types, especially in stem cells that have a high potency, such as embryonic stem cells or induced pluripotent stem cells.

## Acknowledgments

We thank all members of the Canman and Shirasu-Hiza labs for their feedback, support, and advice on this work. We thank Adriana Hernandez and Michelle (Mimi) Schmidt for making worm plates and other critical lab reagents. We thank Mohan Balasubramanian and Geraldine Seydoux for helpful discussions. We are grateful to Drs. Jessica Feldman, Geraldine Seydoux, Amy Maddox, Karen Oegema, and the *Caenorhabditis* Genetics Center (NIH P40OD010440) for providing worm strains. This work was funded by: NIH R01GM117407 (JCC), NSF DGE- 2036197 (JTW), NIH R35GM138380 (KEK), a Sloan Research Fellowship (KEK), a Packard Fellowship (KEK), European Research Council CoG ChromoSOMe N°819179 (JD), NIH R01AG045842 (MSH), and NIH R35GM127049 (MSH). The authors declare no competing financial interests.

## Author contributions

C.Q. Connors, T.R. Davies, M.S. Mauro, J.T. Wiles, and J.C. Canman conceived of the project and designed all experiments. C.Q. Connors, J.T. Wiles, and T. Davies did the mini-RNAi screen to identify genes required for P2 cytokinesis in *formin(ts)* embryos. M.S. Mauro performed all injections for injection-based RNAi depletion (except four replicates among the PIE-1 and POS-1 RNAi validation experiments, which were performed by J.T. Wiles) and performed all quantitative imaging and analysis of contractile ring protein levels. C.Q. Connors conducted all imaging and analysis of cytokinesis outcome experiments, contact angles in 4-to-8-cell embryos, central spindle assembly kinetics, and imaging of fluorescently-tagged PIE-1, POS-1, and septin^UNC-59^ levels with and without double RNAi (fluorescent intensity analysis done by M.S. Mauro), except for single cell cytokinesis outcome analysis upon CCCH Zn-finger depletion in control embryos, which was done by S.L. Martin, or in *formin(ts)* embryos without RNAi treatment or with *anillin^ani-1^(RNAi)*, which was done by J.T. Wiles. J.T. Wiles imaged and quantified fluorescently-tagged CCCH Zn-finger protein levels with and without RNAi in control embryos and performed the P2 spindle, daughter cell size, and microtubule growth rate imaging and analysis. A.D. Countryman and K.E. Kasza did the cell tension data analysis (based on cell-cell contact angles). C.Q. Connors, M.S. Mauro, J.T. Wiles, A.D. Countryman, K.E. Kasza, B. Lacroix, M. Shirasu-Hiza, J. Dumont, and T.R. Davies, and J.C. Canman made significant intellectual contributions and helped write (or edit) the manuscript. C.Q. Connors, M.S. Mauro, J.T. Wiles, A.D. Countryman, and J.C. Canman made the figures.

## Methods

### Worm husbandry and strain maintenance

The *C. elegans* strains used in this study can be found in **Table S1**. Strains were maintained on standard non-vented 60 mm plates (T3308, Tritech Research) filled (PourBoy 4, Tritech research) with 10.5 mL nematode growth media (NGM) (23 g Nematode Growth Medium (Legacy Biologicals, a division of Research Products International (RPI)), 1 mL 1M CaCl_2_, 1 mL of 1M MgSO_4_ 25 mL of 1M K_3_PO_4,_ 975 mL ddH_2_O) seeded with 500 μL OP50 *E. coli* bacteria as a food source, similar to as described^134^. Strains were maintained at 16°C (temperature sensitive mutants) or at 20°C (all other strains) in heating/cooling incubators (Binder).

### RNA-mediated interference (RNAi)

#### Feeding RNAi (mini-screen)

For the mini-screen to identify genes required for protection of P2 cytokinesis in *formin(ts)* embryos (**Figure 1C**), we used feeding RNAi to knockdown candidate genes either implicated in germ fate regulation in the literature or differentially expressed in the P2 cell (relative to the ABa cell) by single cell transcriptomics^96^. Briefly, ∼1000 bp of sequence from the desired gene was amplified by PCR (from cDNA), cloned into the L4440 vector using standard cloning techniques, and transformed into HT115 *E. coli* using CaCl_2_ transformation, as described^135^. RNAi feeding bacteria were grown in LB liquid cultures with 100 μg/mL ampicillin at 32°C for ∼16 hours and 300 μL of each culture was plated on an individual 60 mm RNAi plate (standard NGM plus 50 μg/mL ampicillin and 1 mM IPTG). HT115 *E. coli* with the empty L4440 vector was used as a feeding RNAi control. RNAi plates were allowed to dry and grow at room temperature for 48 hours. *Formin(ts)* L1 larvae were then plated onto RNAi plates and placed in the 16°C incubator for 6 days to become gravid adults before they were dissected to obtain embryos. RNAi primers and template DNA for each target gene are listed in **Table S1**.

#### Injection RNAi (all other experiments)

For all other experiments in the manuscript, we used injection RNAi, which in our hands is more robust than feeding RNAi. Briefly, ∼1000 bp of sequence from the desired gene was amplified by PCR (from cDNA and within a single exon when possible) using primers containing a T7 sequence, confirmed on a 1% agarose gel, PCR purified (QIAquick PCR Purification kit, QIAGEN), and used in T7 reverse transcription reactions (MEGAscript, Life Technologies). The synthesized dsRNAs were purified using phenol-chloroform. The newly synthesized dsRNA was mixed 1:1 with phenol-chloroform (Invitrogen) and mixed by vortexing for 2 minutes. The dsRNA was spun down for 3 minutes at 12,000 x g. The aqueous layer was transferred to new tube, mixed again 1:1 with phenol-chloroform, and vortexed a second time for 2 minutes. The dsRNA was spun down for 3 minutes at 12,000 x g a second time. The aqueous layer was transferred to a new tube and then mixed 1:1 with pre-chilled (-20°C) isopropanol (100%, Sigma) and incubated at -20°C overnight. The dsRNA was precipitated by spinning the sample down at 12,000 x g for 15 minutes. The pellet was allowed to air dry for 5 minutes and then resuspended in 1x soaking buffer (32.7 mM Na_2_HPO_4_, 16.5 mM KH_2_PO_4_, 6.3 mM NaCl, 14.2 mM NH_4_Cl). RNA reactions were annealed at 68°C for 10 minutes followed by 37°C for 30 minutes. dsRNAs were brought to a final concentration of ∼2000-2500 ng/μL (when possible) and 2 μL aliquots of the dsRNA were stored at -80°C until use. For each experiment, a fresh aliquot (or aliquots for double RNAi experiments) was diluted to ∼1000 ng/μL (∼500 ng/μL for *pie-1* and *pos-1* dsRNA and ∼1000 ng/μL for *septin^unc-59^* dsRNA in the double RNAi experiments) using 1x soaking buffer and centrifuged at 13,000 rpm for 10 minutes at room temp (∼22°C). 0.35 μL of the diluted dsRNA was loaded into the back of pulled borosilicate glass capillary needles (World Precision Instruments, WPI; Sutter Instruments, P1000 needle puller) and injected into the gut of L4 worms using a Leica DMIRB microscope equipped with Hoffman optics, a Plan L 20x/0.4 CORR PH (Leica), a rotating stage, and the XenoWorks digital microinjector and micromanipulator injection system (Sutter Instruments). Worms were rescued by resuspension in M9 buffer (6 g KH_2_PO_4_, 12 g Na_2_HPO_4_, 10 g NaCl, 0.5 mL 1 M MgSO_4_, ddH_2_O to 2 L) to plates seeded with OP50 bacteria and allowed to recover for ∼24 hours at 20°C or ∼42 hours at 16°C (temperature sensitive strains) prior to imaging or embryonic lethality analysis. RNAi primers and template DNA for each target gene are listed in **Table S1**.

### Embryonic lethality analysis

#### Embryonic lethality quantifications for the mini-screen (feeding RNAi; Figure S1B)

On the morning of each experiment, 5 young adult/adult hermaphrodite worms from both the experimental group (*formin(ts)* plus candidate gene targeting-RNAi) and the control group (*formin(ts)* plus control RNAi (empty L4440 vector)) were singled out onto non-vented 35 mm NGM plates (T3501, Tritech Research) with 100 µL of OP50 and placed back into the 16°C incubator (permissive temperature for the *formin(ts)* worms). Hermaphrodites were allowed to lay eggs for the duration of the day. In the evening (7-10 hours later), the adult worm was removed from each plate. The following day, each plate was manually scored for hatched larvae and unhatched embryos on a high-resolution dissecting microscope (Olympus SZX16 with an Olympus SDF PLAPO 1XPF objective).

#### Embryonic lethality quantifications for injection RNAi

L4 hermaphrodite worms were injected with indicated dsRNA and allowed to recover for 42 hours at 16°C. 42-hours post-injection, dsRNA-injected and control adult worms were then singled out and allowed to self-fertilize for 24 hours, then the adult worms were disposed of. Prior to counting, embryos were given 36 hours at 16°C to hatch. Plates were scored for hatched larvae and unhatched (dead) embryos on a high-resolution dissecting microscope, as above (feeding RNAi).

### Embryo preparation for live cell imaging

Young gravid adult hermaphrodites were kept at 13-14°C in a small incubator (Wine Enthusiast, model 2720213W) dissected on a high-resolution dissecting microscope (Olympus SZX16 with an Olympus SDF PLAPO 1XPF objective) in cooled (13-14°C) M9 buffer. 4-cell stage embryos were mounted on a thin (∼1-2x lab tape thickness) 2% agar pad on a glass slide (VWR VistaVision, 3 inches x 1 inch x 1 mm) using a hand-pulled glass pipette (VWR Pasteur Pipette) or a borosilicate glass capillary (World Precision Instruments, WPI) as a mouth pipette. A 22 x 22 mm No. 1.5 glass coverslip (VWR) was placed on top of the embryos for imaging, similar to as described^136^.

### Live cell imaging set up and temperature control

We used two microscopes for live imaging experiments. Both microscope systems were controlled by MetaMorph software (Molecular Devices). Live imaging was performed in an imaging room equipped with a heat-pump based temperature control device (see below for details of each system). Room and microscope temperatures were continuously monitored using 4-5 digital thermometers placed around the room and near the microscope stage and a Bluetooth-enabled smart temperature sensor (SensorPush) on the microscope stage. All imaging was done at 26 +/- 0.5°C, except for in **Figure 2** and **Figure 8** (and **Figure 3, Figure S3**) which was done specifically at 25.5-26.0°C.

### Live imaging microscopes

For the P2 cytokinesis RNAi mini-screen (**Figure 1C**), single cell cytokinesis outcome experiments (**Figure 2**), central spindle assembly analysis (**Figure 4E-F, Figure S5**), half of the surface tension experiments (using the same *formin(ts)* control and *formin(ts); pie-1(RNAi)* data shown in **Figure 2B,C** and presented in **Figure 3, Figure S3**), and quantitative analysis of contractile ring protein levels (**Figure 5, Figure S6, Figure 6, Figure S7, Figure 7, Figure S9, Figure S10**), live cell imaging was performed in an imaging room with a mini-split heat pump-based temperature control device (MultiAqua; model MHWX Hi-Wall Fan Coils). The microscope was built on an inverted stand (Nikon, Eclipse Ti) with a spinning disc confocal (Yokogawa, CSU-10 with Borealis (Spectral Applied Research)), a charge-coupled device (CCD) camera (Hamamatsu Photonics, Orca-R2), and a Piezo-driven motorized stage (Applied Scientific Instrumentation, ASI) for Z-sectioning. Focus was maintained (Nikon, Perfect Focus) before each Z-series acquisition. Excitation laser light (150 mW 488 nm and 561 nm, Spectral Applied Research (ILE-2)) was controlled by an acousto-optic tunable filter (Spectral Applied Research), and a filter wheel (Sutter Instruments) was used for DIC analyzer and emission filter (525/50 nm and 620/60 nm bandpass (Chroma)) selection.

For quantitative analysis of protein levels after RNAi treatment (**Figure S2A-D, Figure S8**), the other half of the surface tension experiments (using the same *formin(ts)* control and *formin(ts); pie-1(RNAi)* data shown in **Figure 8**, and presented in **Figure 3, Figure S3**), spindle dynamics analysis (**Figure 4A-C, Figure S4**), microtubule growth rate analysis (**Figure 4D**), and some P2 cytokinesis outcome experiments (**Figure 8**), live cell imaging was performed in an imaging room with a mini-split heat pump-based temperature control device (Mitsubishi; Mr. Slim, MSZ-D36NA). The microscope was built on an inverted stand (Nikon, Eclipse Ti; custom-modified for compatibility with near-infrared light, as in^137,138^) with a spinning disc confocal (Yokogawa, CSU-10 with Borealis (Spectral Applied Research)), a CCD camera (Hamamatsu Photonics, Orca-R2), and a Piezo-driven motorized stage (ASI) for Z-sectioning. Focus was maintained (ASI, CRISP) before each Z-series acquisition. Two solid state 150 mW 488 nm and 561 nm lasers (Cairn) were used for excitation light, and a filter wheel (Ludl Instruments) was used for DIC polarizer and emission filter (525/50 nm and 620/50 nm bandpass (Chroma)) selection.

### Live cell imaging and analysis parameters

#### FIJI (FIJI is Just ImageJ) software^139^ was used for all data analyses

***Single cell cytokinesis outcome analysis:*** For the P2 cell cytokinesis mini-screen (**Figure 1C**) we used a 20x Plan Apo 0.75 N.A. dry objective (Nikon) with 2 x 2 binning and 13 x 2 µm Z-sections every 60 seconds. For all other single cell cytokinesis outcome analysis experiments and for half of the surface tension measurements (using the same data sets shown in **Figure 8**, shown in **Figure 3, Figure S3**) we used a 60x Plan Apo 1.40 N.A. oil immersion objective (Nikon) with 2 x 2 binning and 15 x 2 µm Z-sections every 60 seconds. Cytokinesis outcome was scored manually by eye on maximum projection images of both channels (GFP::PH^pLCδ^ and mCherry::histone H2B^HIS-58^)^97^. Individual cells were only scored if the image series began before anaphase onset and ended after at least one of that cell’s daughter cells entered anaphase of the next cell cycle, except for RNAi experiments in control embryos (**Figure 2A**), in which embryo viability was assumed. In those control embryos (**Figure 2A**), completion was scored if the image series began before anaphase onset and ended after a dividing membrane visible across all Z planes persisted between the cell’s daughter cells for at least 180 seconds. In all other cytokinesis outcome analyses, cytokinesis in each individual cell was scored as either completed successfully (the cell under observation divided into two daughter cells and the contractile ring remained closed when a daughter cell entered anaphase of the next cell cycle) or failed (little to no contractile ring constriction or partial or full cleavage furrow ingression followed by contractile ring regression and binucleation).

#### Surface tension analysis

For P2 and neighboring cell surface tension analysis (**Figure 3, Figure S3**) we used a 60x Plan Apo 1.40 N.A. oil immersion objective (Nikon) with 2 x 2 binning and 15 x 2 µm Z-sections every 60 seconds. The data set used for this analysis came from two data sets used for analyses elsewhere in this paper: data collected for P2 cytokinesis outcome experiments shown in **Figure 2B,C** and **Figure 8**. Image analysis was done on a single Z plane of the 4-cell embryo that was determined to be the most central to the longest and widest aspects of the P2 cell based on the fluorescently-tagged plasma membrane reporter. The time point at which the P2 cleavage furrow was first visible was considered t=0 s (P2 furrow onset) and the previous 6 time points were also included in the analysis. At each time point, the FIJI^139^ angle tool was used to measure the three different membrane contact angles at each of the two locations that the P2 cell forms a cell-cell contact with a neighboring cell (either the ABp cell on the dorsal side or the EMS cell on the ventral side).

Cell surface tension analysis was done as recently described^101^ with a few modifications. Measured angles were re-scaled to add to 360° for self-consistency. A Young-Dupré force balance was used to relate P2 surface tension to EMS/ABp surface and interfacial tensions using the measured contact angles:

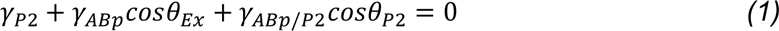

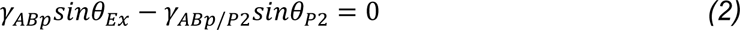

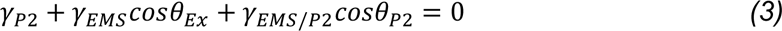

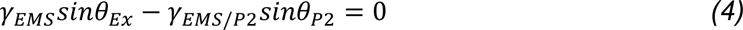

#### Quantitative analysis of fluorescently-tagged reporters for RNAi knockdown efficiency

For quantitative imaging of fluorescently-tagged MEX-1^98^, PIE-1^99^, POS-1^100^, and septin^UNC-59^ ^117^ reporters with and without the respective single or double dsRNA injection (**Figure S2A-D and Figure S8**), we used a 60x Plan Apo 1.40 N.A. oil immersion objective (Nikon) with 2 x 2 binning and 15 x 2 µm Z-sections. Image analysis was done on sum projection images of 4-cell stage embryos from each individual fluorescently-tagged CCCH Zn-finger protein or septin^UNC-59^ reporter strain with and without RNAi-mediated depletion of that Zn-finger protein or septin^UNC-59^. A region surrounding the P2 cell (for Zn-finger proteins) or whole embryo (for *septin^unc-59^* knockdown) was manually drawn to measure the total fluorescence intensity and a 20 x 20-pixel box was drawn in the cytoplasm of the ABa cell to calculate intracellular embryonic background levels. The average ABa cytoplasmic fluorescence background intensity was multiplied by the measured area of the P2 cell (for Zn-finger proteins) or whole embryo (for *septin^unc-59^* knockdown) area and then subtracted from each P2 cell (or whole embryo) fluorescence intensity measurement.

#### P2 spindle and cellular dynamics imaging and analysis

For analysis of P2 spindle and cellular dynamics with and without *pie-1(RNAi)* (**Figure 4A-C, Figure S4**), we used a 60x Plan Apo 1.40 N.A. oil immersion objective (Nikon) with 1 x 1 binning and 13 x 1.5 µm Z-sections every 20 seconds. This analysis was done in a strain expressing mCherry::histone H2B^97^ to label the chromosomes, mCherry::PH^pLCδ 140^ to label the plasma membrane, and EB1^EBP-2^::GFP^104^ to label the centrosomes. All measurements were taken using the line drawing tool in FIJI^139^. Composite images were made of both mCherry and GFP channels at all Z planes and measurements were taken on the Z plane where both objects of interest (chromosomes, centrosomes, or polar cell cortex plasma membrane) were best visible and in focus (*e.g.*, the Z plane containing the P3- destined centrosome and chromosomes or the C-destined centrosome and chromosomes). Forming daughter cell size measurements were taken on the Z plane in which the distance between the cell cortex plasma membrane was the greatest. For the centrosome to centrosome, chromatin to chromatin, and P2 cell long axis measurements specifically, maximum intensity projections were used to enable visualization of objects in different Z planes.

#### Microtubule growth rate imaging and analysis

For analysis of astral microtubule growth rates with and without *pie-1(RNAi)* (**Figure 4D**), we used a 60x Plan Apo 1.40 N.A. oil immersion objective (Nikon) with 1 x 1 binning and a single Z-plane every 0.5 seconds for at least 2 minutes. Embryos were monitored by DIC and mCherry::histone H2B^HIS-58^ ^97^ until anaphase onset, at which point time lapse imaging was initiated. Image analysis was done using the “Manual Tracking” plugin in FIJI^139^. The plugin was used to measure velocities of microtubule plus tips that could be clearly seen for three frames, based on the distance traveled per frame, the temporal resolution between frames, and the xy-calibration of the microscope. Two velocities for each microtubule plus tip were calculated by clicking on the center of the microtubule plus tip across three frames; the plugin calculated one velocity between the first and second frames and a second velocity between the second and third frames. An average velocity for each microtubule plus tip was then determined by averaging the two velocities calculated with the plugin.

#### Central spindle assembly imaging and analysis

For analysis of central spindle assembly with and without *pie-1(RNAi)* (**Figure 4E-F, Figure S5**), we used a 40x Plan Fluor 1.30 N.A. oil immersion objective (Nikon) with 2 x 2 binning and 10 x 1.25 µm Z-sections every 10 seconds. Embryos were monitored by DIC and mCherry::histone H2B^107^ until the initiation of nuclear envelope breakdown, at which point Aurora-B^AIR-2^::GFP^105^ imaging was started. Quantitative measurements of peak Aurora-B^AIR-2^::GFP and mCherry::histone H2B levels began at the frame prior to anaphase onset (metaphase). For each measurement and timepoint, a linescan of 10 µm wide and 80 µm long was drawn from the “EMS side” to the “ABp side” of the P2 cell, perpendicular to the division plane in both the GFP and mCherry channels. An extracellular background linescan was also taken for each channel. The background linescan for each channel was averaged and then subtracted from each pixel value along the P2 central spindle linescan in each channel.

#### Quantitative analysis of fluorescently-tagged contractile ring protein levels

For quantitative imaging of fluorescently-tagged F-actin reporters (LifeAct::RFP and plastin^PLST-1^::GFP^115^), myosin-II^NMY-2^::GFP^116^, septin^UNC-59^::GFP^117^, and anillin^ANI-1^::GFP^118^) with and without dsRNA injection in control and *formin(ts)* embryos (**Figures 5-7, Figure S6, Figure S7, Figure S9, Figure S10**), we used a 60x Plan Apo 1.40 N.A. oil immersion objective (Nikon) with 2 x 2 binning and a single 45 x 0.5 µm Z-section timepoint was taken. Embryos were monitored by DIC and images were acquired at the time of cleavage furrow onset in the P2 cell (LifeAct::RFP, plastin^PLST-1^::GFP, myosin-II^NMY-2^::GFP, and anillin^ANI-1^::GFP) or 20s after P2 furrow onset (septin^UNC-59^::GFP). For image analysis, first embryos were rotated so the P2 division plane was located on the right side of the image. Next sum projections were created for each embryo. A 25 x 50-pixel box was drawn over the P2 cleavage furrow/division plane and the total fluorescence intensity was measured. To measure the total fluorescence intensity of the entire embryo, an oval was drawn around the embryo. A 25 x 50-pixel box was then placed outside of the embryo to measure the extracellular background (camera background) and value was then subtracted from the total fluorescence intensity in the division plane. For the whole embryo total fluorescence intensity, background was subtracted by multiplying the mean intensity of the extracellular background by the area of the embryo. Each individual value was than normalized to the control average for its respective experimental replicate. For **Figure 7** individual data points are shown and normalized to the *formin(ts)* average.

To measure the asymmetric localization of myosin-II^NMY-2^::GFP, septin^UNC-59^::GFP, and anillin^ANI-1^::GFP on the dividing P2 cell cortex, two independent 25-pixel wide x 10 µm long lines were drawn across the future C-cell cortex and the future P3-cell cortex from outside of the cell into cytoplasm (see schematic in **Figure 5B**). The first 10 pixels of each linescan from outside of the cell were averaged to calculate the average extracellular background (camera background). The average extracellular background was then subtracted from all other points along the same line. The last 15 pixels of the line (∼2.4 microns; inside the cell) were averaged to calculate the “average cytoplasmic value” for each forming daughter cell. The maximum value at the cell cortex was used as the “peak cortical level”. The peak cortical level measurement was divided by the average cytoplasmic level to calculate the cell cortex to cytoplasmic ratio for each forming daughter cell. The peak cortical value for the forming C cell was divided by the peak cortical value for the forming P3 cell to calculate the ratio between the two. The C-destined cytoplasmic value was divided by the P3-destined cytoplasmic value to calculate the ratio between the two. To plot the linescan data on one graph, each point along the line was normalized using the respective cytoplasmic value.

### Figure preparation

All figures were made using Adobe Illustrator 2023; graphs were made in Prism 9 (Graphpad) or in Python using Matplotlib^141^ (tension analysis only) and pasted into Adobe Illustrator.

### Statistical analysis

All statistical tests were done in Prism 9 (Graphpad) or by using Excel (Microsoft). For all single cell cytokinesis outcome experiments, a Fisher’s exact test was used. For analysis of significance in most experiments an unpaired Student’s t-test was used (with a Holm-Sidak correction for surface tension analysis only) except for the quantitative analysis of mean microtubule growth rates and anillin^ANI-1^ levels at the P2 division plane where a two-way ANOVA was used. Error bars in all graphs represent the standard deviation (SD). p-values: n.s.=p≥0.05, *=p<0.05, **=p<0.01, ***=p<0.001, and ****=p<0.0001; except for tension analysis (n.s.=p-value not significant; *=p-value significant; Figure 3**, Figure S3**); see also **Table S1**.

## Supplemental Figure Legends

**Figure S1:**
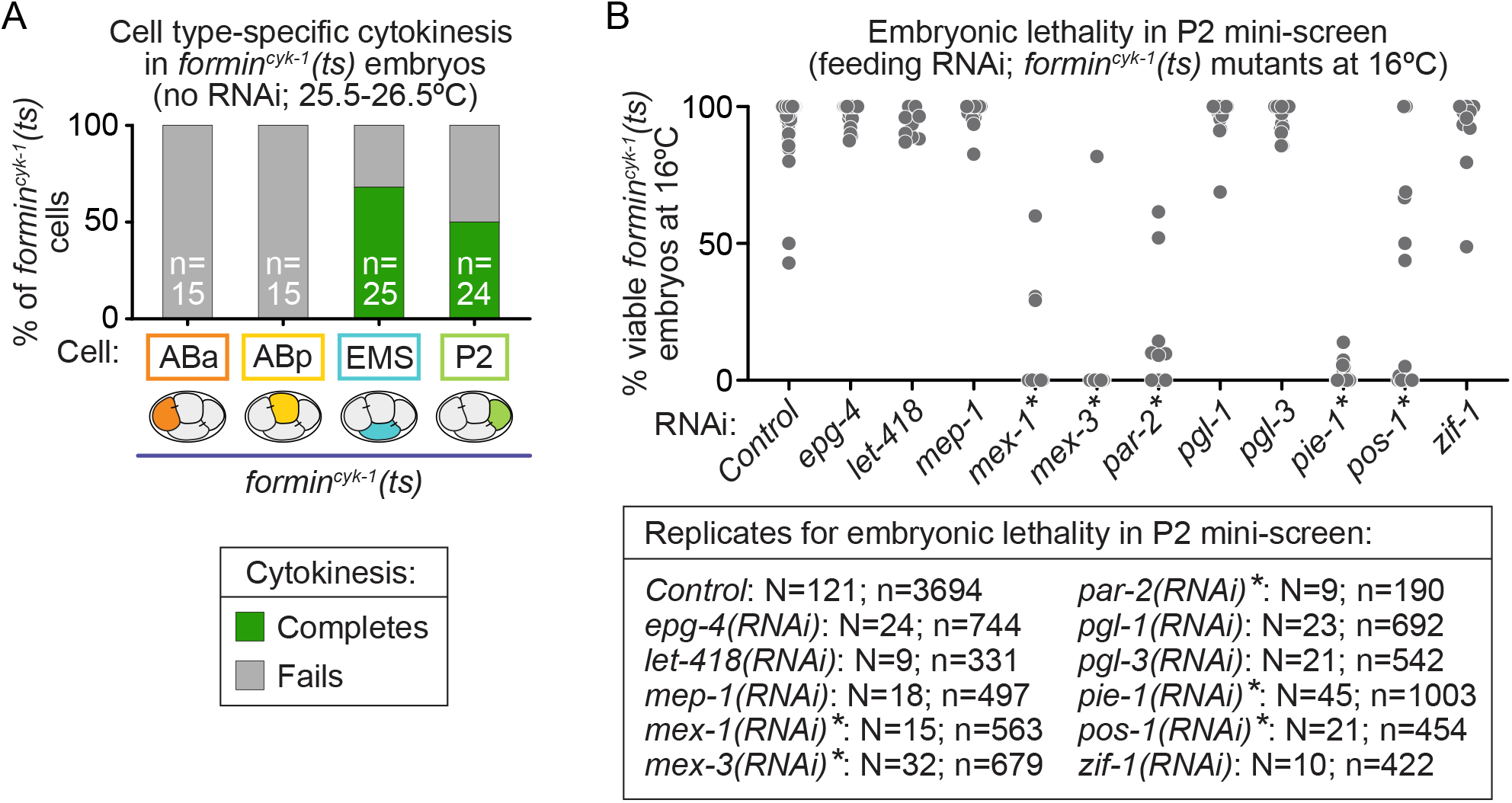
Cell type-specific cytokinesis protection in formin^cyk-1^(ts) embryos and embryonic lethality from RNAi-based mini-screen. **A)** Graph showing the percentage of ABa, ABp, EMS, and P2 cells that complete (green) or fail (gray) cytokinesis in 4-to-8-cell *formin^cyk-1^(ts)* embryos at restrictive temperature; n=number of individual cells scored and is indicated on each bar. **B)** Graph plotting the percentage of viable embryos after each indicated feeding RNAi treatment (related to Figure 1C); *=RNAi-knockdown of candidate gene expected to result in embryonic lethality^142^; N=number of worms; n=number of progeny (embryos and larvae).

**Figure S2:**
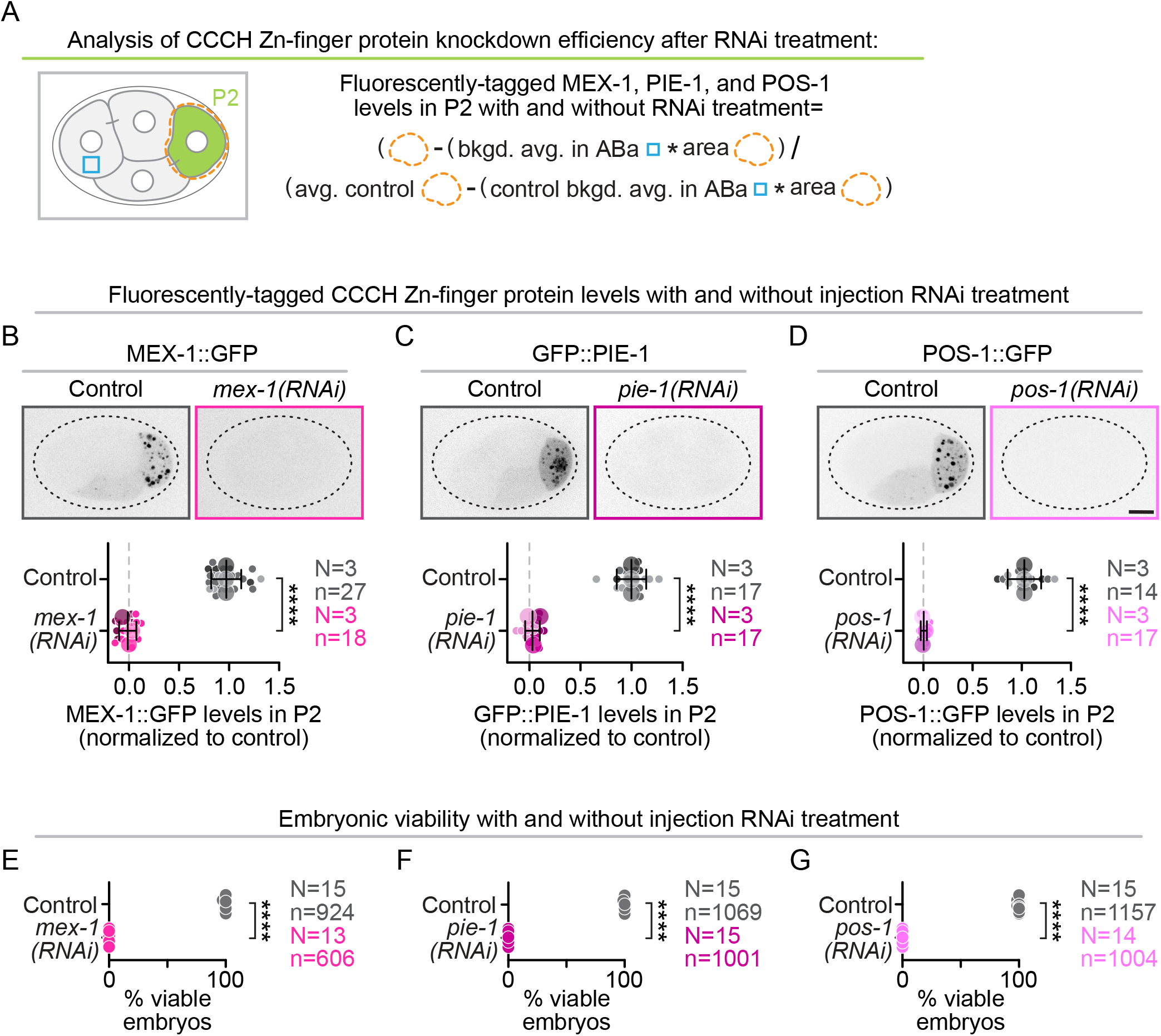
Injection RNAi knockdown efficiency of CCCH Zn-finger proteins. **A)** Schematic depicting analysis shown in **B-D** performed on sum projected images to measure CCCH Zn-finger protein levels in the P2 cell (dashed orange line) and embryonic background (blue box in ABa cell, see also Methods) with and without injection RNAi treatment. **B-D)** Representative maximum projection inverted contrast images (top panels) and graphs (bottom panels) showing quantification (from sum projected images) of the efficiency of MEX-1, PIE-1, and POS-1 knockdown after each RNAi treatment in 4-cell stage embryos expressing fluorescently tagged **B)** MEX-1::GFP, **C)** GFP::PIE-1, and **D)** POS-1::GFP reporters with and without the corresponding injection RNAi treatment (controls=gray, *mex-1(RNAi)*=fuchsia pink, *pie-1(RNAi)*=raspberry pink, *pos-1(RNAi)*=bubblegum pink). Scale bar=10 μm; error bars=standard deviation; N=number of experimental replicates; n=number of embryos scored for each genotype by color; ****=p-value ≤0.0001 (Student’s t-test, unpaired, see also Table S1). **E-G)** Graphs plotting the percentage of viable embryos from untreated control (gray) and **E)** *mex-1(RNAi)* (fuchsia pink), **F)** *pie-1(RNAi)* (raspberry pink), and **G)** *pos-1(RNAi)* (bubblegum pink) worms. Error bars=standard deviation; N=number of worms; n=number of total progeny (embryos and larvae) for each genotype by color; ****=p-value ≤0.0001 (Student’s t-test, unpaired, see also **Table S1**).

**Figure S3:**
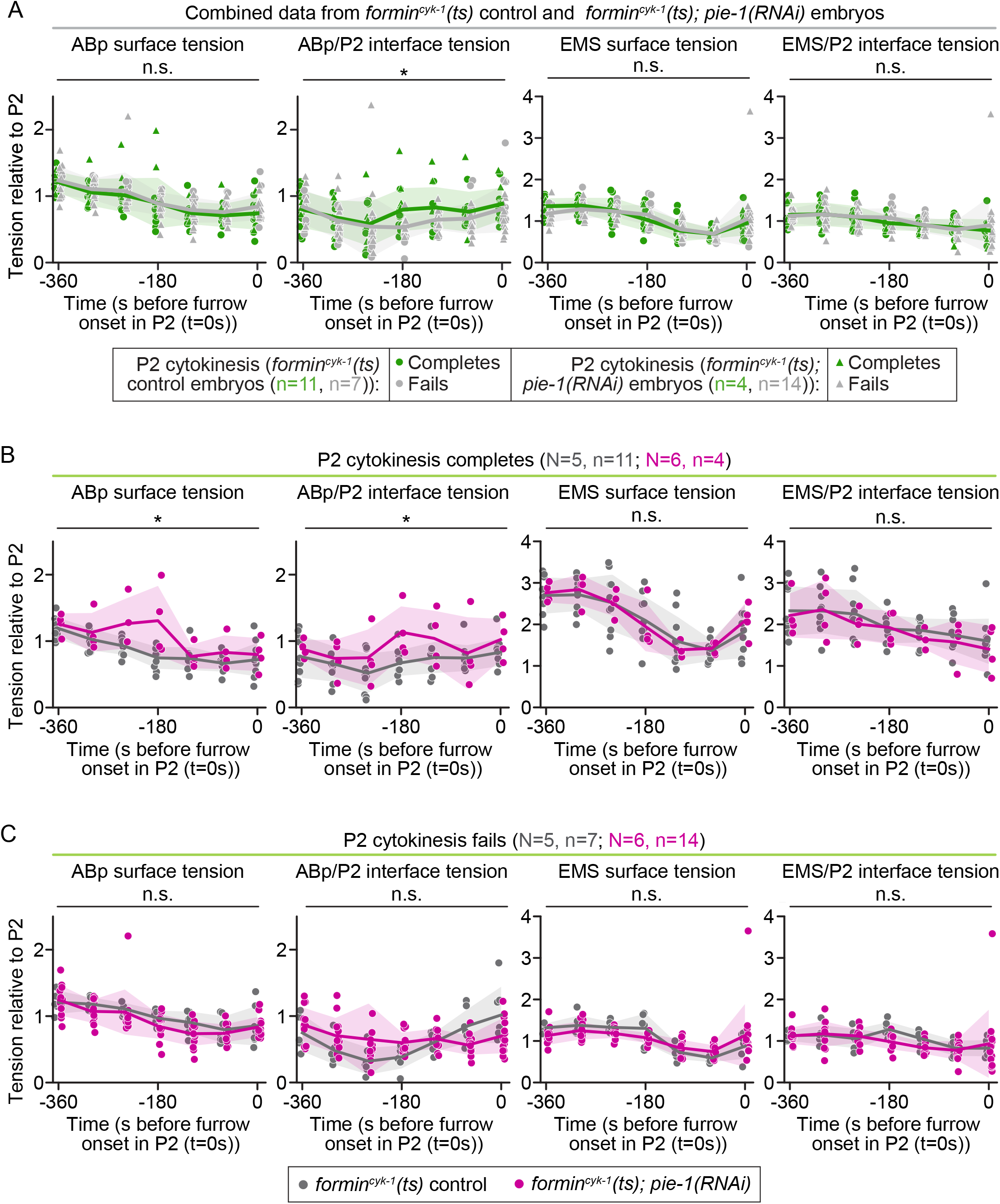
Cytokinesis failure is associated with altered surface tension patterns, but the precise surface tension patterns associated with successful cytokinesis are distinct in control and pie-1(RNAi) embryos. Graphs showing the relative ABp surface tension (far left panels), ABp/P2 interface tension (center left panels), EMS surface tension (center right panels), and EMS/P2 interface tension (far right panels) in **A)** combined *formin^cyk-1^(ts)* control (circles) and *formin^cyk-1^(ts); pie-1(RNAi)* (triangles) embryos in which P2 cytokinesis completes (2 mononucleated daughter cells, green) or fails (1 binucleated daughter cell, gray). Surface tension patterns in *formin^cyk-1^(ts)* control (gray) and *formin^cyk-1^(ts); pie-1(RNAi)* (pink) embryos in which P2 cytokinesis **B)** completes or **C)** fails. Tension measurements normalized to P2 surface tension; time (s) is relative to furrow onset (t=0s) in each P2 cell; tension measurements taken on *formin ^cyk-1^(ts)* control and *formin ^cyk-1^(ts); pie-1(RNAi)* datasets in Figure 2B and Figure 8; see also Methods; N=number of experimental replicates; n=number of embryos scored for each cytokinesis outcome **(A)** or genotype **(B, C)** by color; n.s.=p-value not significant, *=p-value significant (Student’s t-test, unpaired, Holm-Sidak correction; see also **Table S1**).

**Figure S4:**
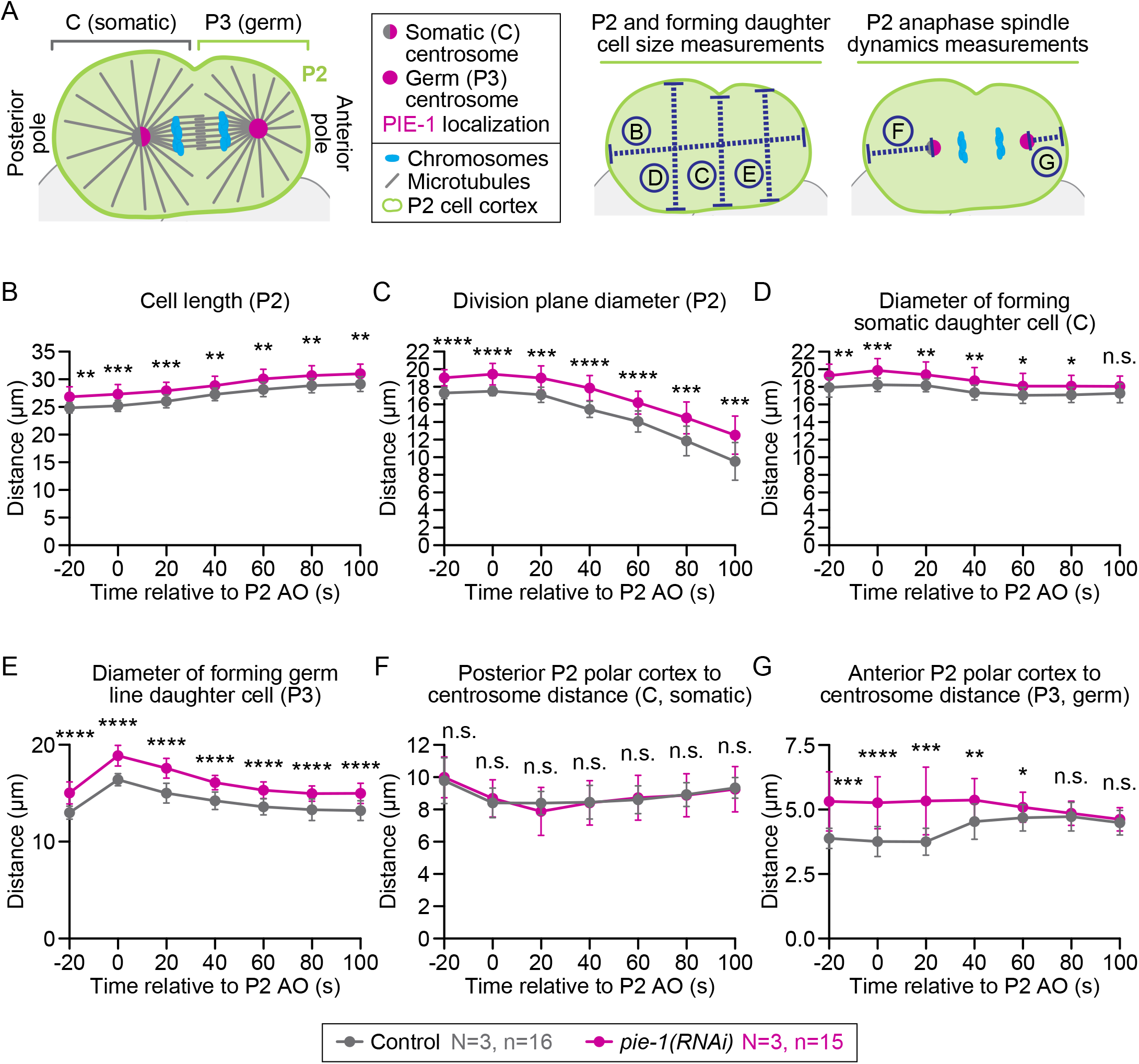
PIE-1 depletion does not affect P2 spindle dynamics and has a minor effect on daughter cell dynamics. **A)** Schematic (left) of the P2 cell during anaphase indicating the P2 cell cortex (lime green), chromosomes (blue), centrosomes (gray/pink for C-destined centrosome, pink for P3-destined centrosome), and microtubules (gray). (Right) Cartoon schematic of measurements taken and plotted in **B-G.** Graphs plotting **B)** P2 cell length, **C)** diameter of the P2 division plane, **D)** diameter of the forming C (somatic) daughter cell, **E)** diameter of the forming P3 (germ) daughter cell, **F)** distance from the posterior polar cortex to the C-destined centrosome, and **G)** distance from the anterior polar cortex to the P3-destined centrosome over time in dividing control (gray) and *pie-1(RNAi)* (pink) P2 cells expressing EB1^EBP-2^::GFP, mCherry::histone H2B^HIS-58^, mCherry::PH^pLCδ^. Time (s) is relative to anaphase onset in each P2 cell (AO, t=0s); error bars=standard deviation; N=number of experimental replicates; n=number of P2 cells scored for each genotype by color; n.s.=not significant, *=p-value ≤0.05, **=p-value ≤0.01, ***=p-value ≤0.001, ****=p-value ≤0.0001 (Student’s t-test, unpaired, see also **Table S1**).

**Figure S5:**
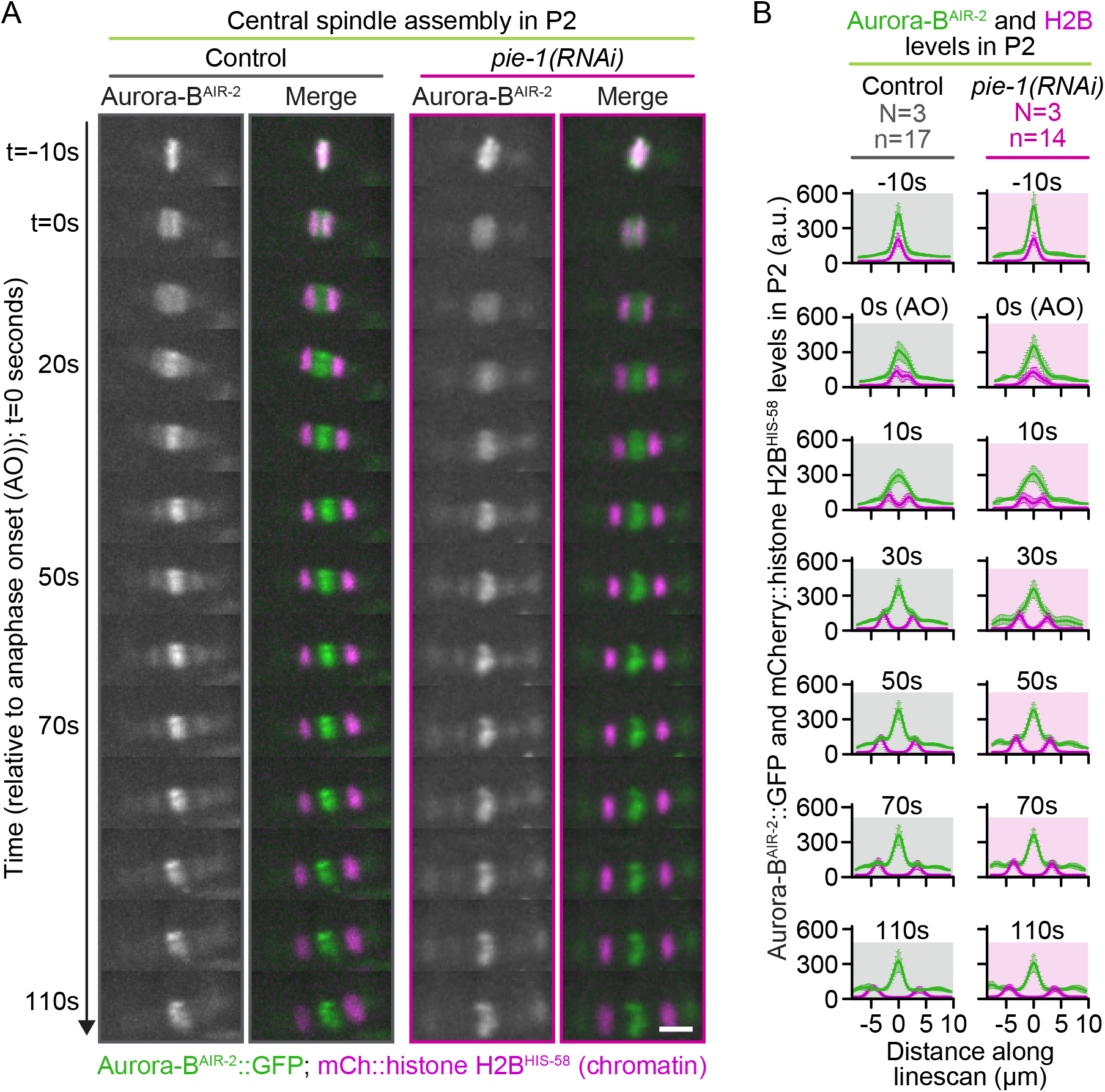
PIE-1 depletion does not affect P2 central spindle assembly. **A)** Representative pseudo-kymographs showing Aurora-B^AIR-2^::GFP (grayscale and green) and mCherry::H2B (magenta) in control (left panels) and *pie-1(RNAi)* (right panels) P2 cells over time. Time on the left; t=0s is anaphase onset in P2; images acquired every 10s; scale bar=10 µm. **B)** Graphs plotting linescan analysis of peak Aurora-B^AIR-2^::GFP (green) and mCherry::H2B (magenta) levels in control (gray, left) and *pie-1(RNAi)* (pink, right) P2 cells over time. Time (s) above each graph; t=0s is anaphase onset (AO) in P2; error bars=standard deviation; N=number of experimental replicates; n=number of embryos scored for each genotype by color (see also Figure 3E-F).

**Figure S6:**
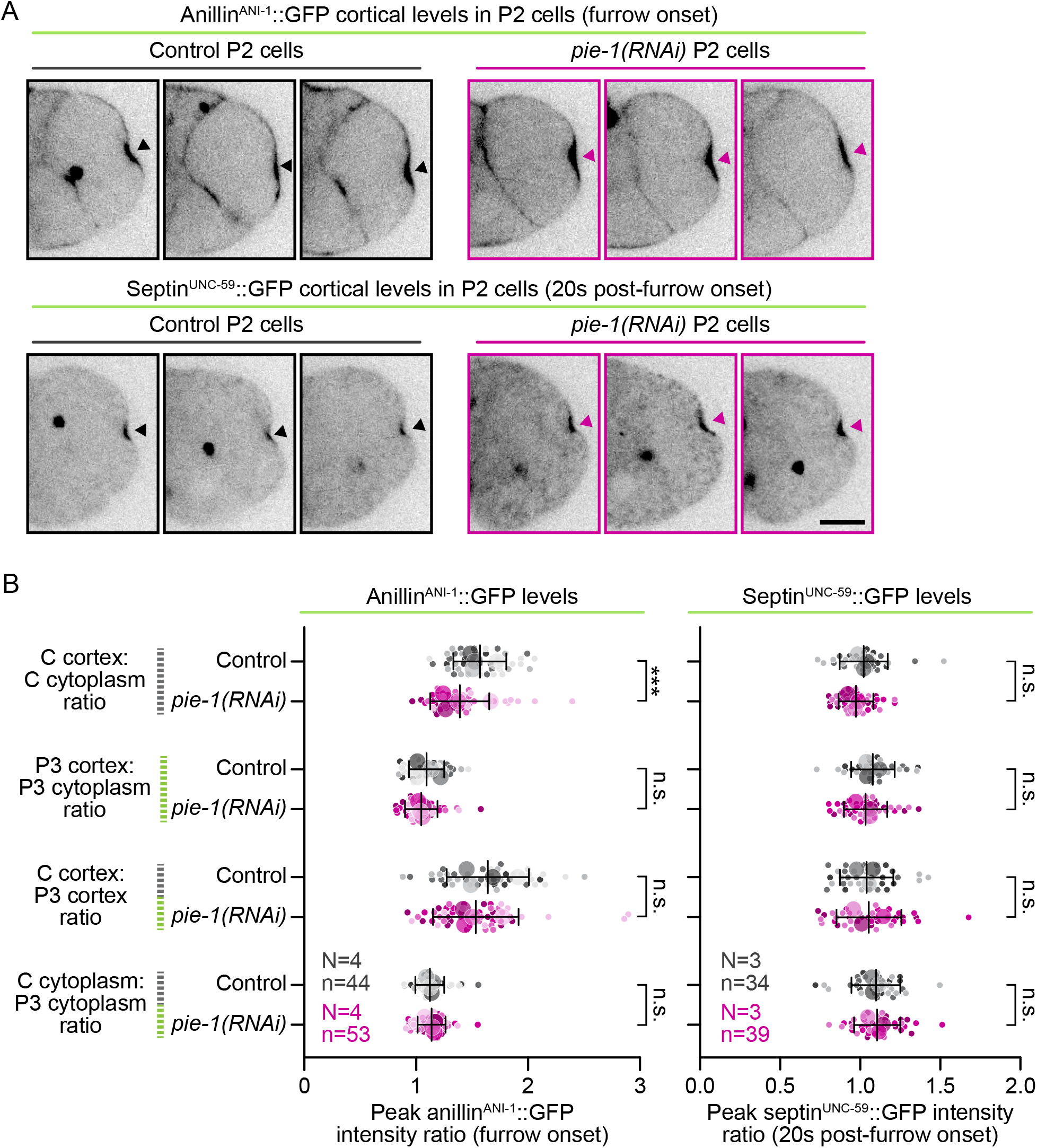
PIE-1 does not play a major role in P2 division asymmetry. **A)** Representative single plane images of 3 wildtype (left) and 3 *pie-1(RNAi)* (right) P2 cells expressing endogenously-tagged anillin^ANI-1^::GFP (top panels) or septin^UNC-59^::GFP (bottom panels) at the time of cleavage furrow onset in P2 (for anillin^ANI-1^::GFP) or 20s post-furrow onset (for septin^UNC-59^::GFP); scale bar=10 μm; gray (control) and pink (*pie-1(RNAi)*) arrowheads indicate the P2 cleavage furrow. **B)** Graph plotting peak cortical intensity ratios for anillin^ANI-1^::GFP (left) and septin^UNC-59^::GFP (right) in control (gray) and *pie-1(RNAi)* (pink) embryos at the forming C and P3 daughter cell cortexes (relative to average cytoplasmic levels) during cell division. Error bars=standard deviation; N=number of experimental replicates; n=number of embryos scored for each genotype by color; n.s.=not significant, ***=p-value ≤0.001 (Student’s t-test, unpaired, see also **Table S1**; see schematic in Figure 5B and Methods).

**Figure S7:**
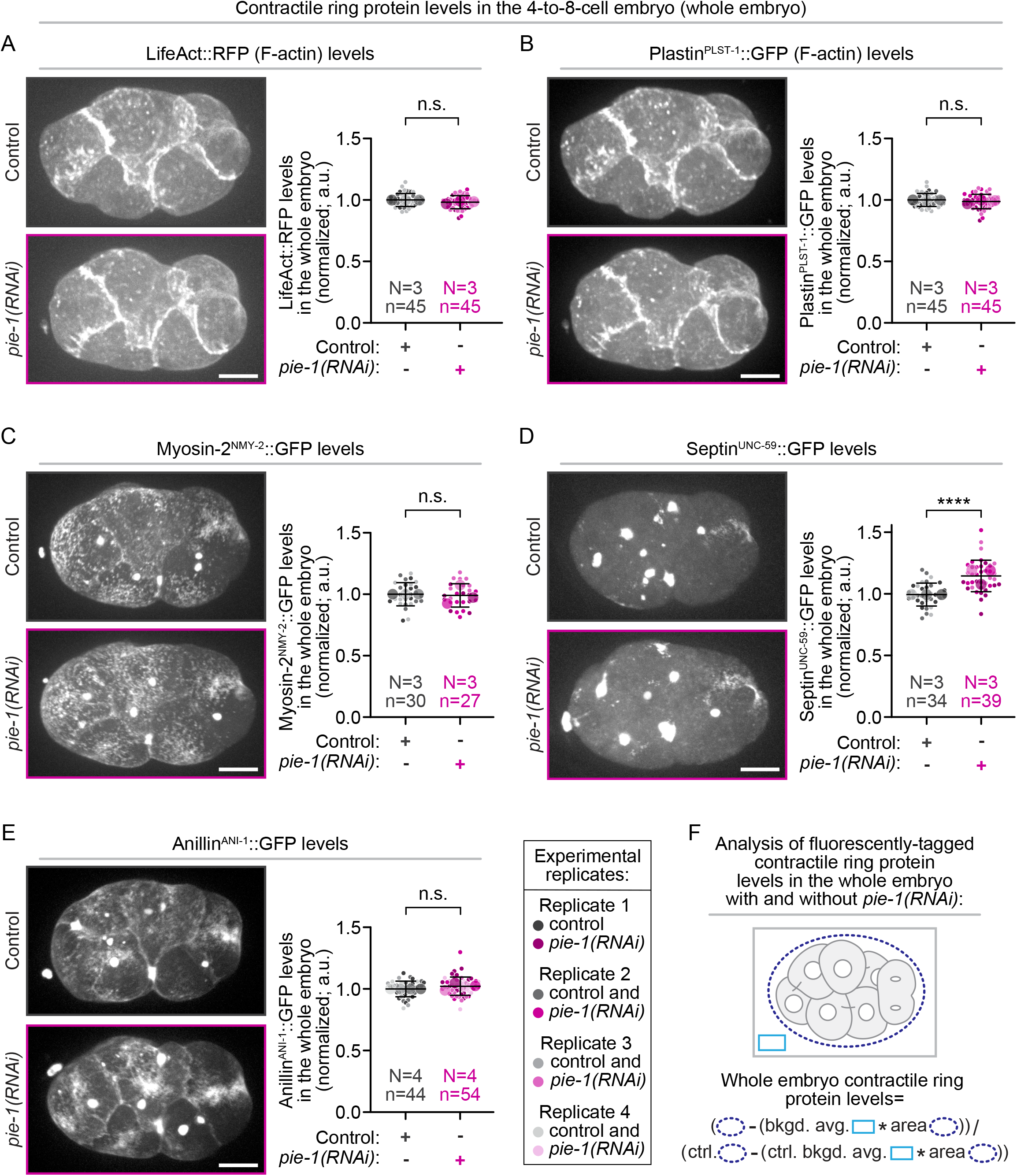
PIE-1 restricts the levels of septin^UNC-59^, but not anillin^ANI-1^, in the 4-to-8-cell embryo. Representative maximum projection images (left) and graphs (right) showing super plots of normalized whole embryo levels of indicated fluorescent reporter for wildtype (grays) and *pie-1(RNAi)* (pinks) embryos expressing fluorescently tagged reporters for **A, B)** F-actin (**A)** Lifeact and **B)** Plastin^PLST-1^), **C)** myosin-II^NMY-2^, **D)** septin^UNC-59^, and **E)** anillin^ANI-1^ at the time of cleavage furrow onset in the P2 cell. Small circles indicate individual data points and large circles and color shades indicate replicate averages; scale bar=10 μm; error bars=standard deviation; N=number of experimental replicates and n=number of embryos scored for each genotype by color; n.s.=not significant, ****=p-value ≤0.0001 (Student’s t-test, unpaired, see also **Table S1**). **F)** Schematic depicting analysis shown in **A-E** performed on sum projected images to measure contractile ring protein levels in the 4-to-8-cell embryo (dark blue dashed oval) and extracellular background (light blue box, see also Methods).

**Figure S8:**
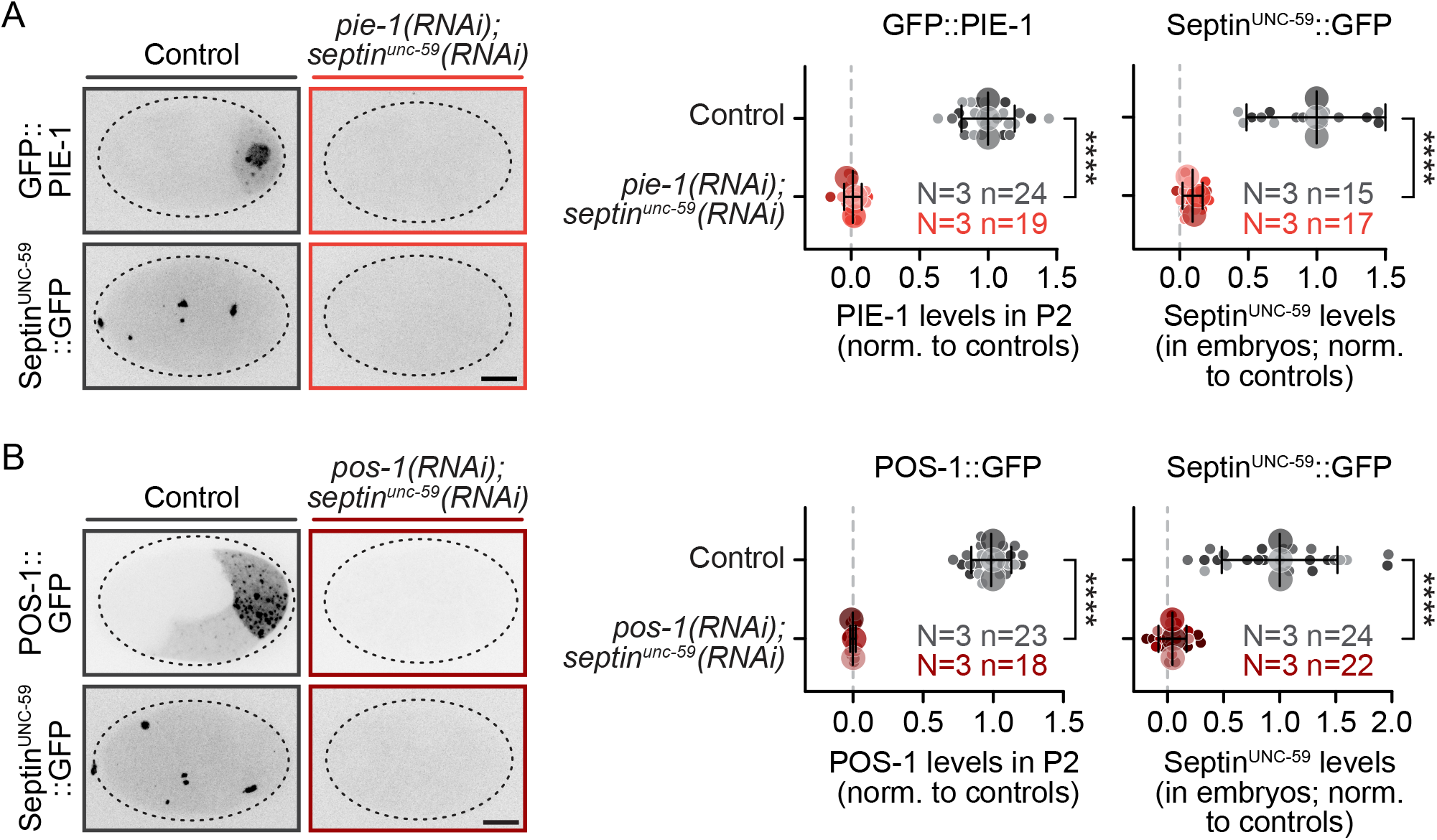
Injection RNAi efficiency for pie-1 or pos-1 and septin^UNC-59^ double knockdowns. Representative maximum projection inverted contrast images (left panels) and graphs (right panels) showing quantification of the efficiency of **A)** *pie-1(RNAi); septin^unc-59^(RNAi)* in 4-cell stage embryos expressing fluorescently tagged GFP::PIE-1 (top panels, see schematic in **Figure S2A**, see also Methods) or septin^UNC-59^ (bottom panels, see schematic in **Figure S7F**, see also Methods) and **B)** *pos-1(RNAi); septin^unc-59^(RNAi)* in 4-cell stage embryos expressing fluorescently tagged POS-1::GFP (top panels, see schematic in **Figure S2A**, see also Methods) or septin^UNC-59^ (bottom panels, see schematic in **Figure S7F**, see also Methods) with and without the corresponding double injection RNAi treatment (controls=grays, *pie-1(RNAi); septin^unc-59^(RNAi)*=reds, *pos-1(RNAi); septin^unc-59^(RNAi)*=maroons). Scale bar=10 μm; error bars=standard deviation; N=number of experimental replicates; n=number of embryos scored for each genotype by color; ****=p-value ≤0.0001 (Student’s t-test, unpaired, see also **Table S1**).

**Figure S9:**
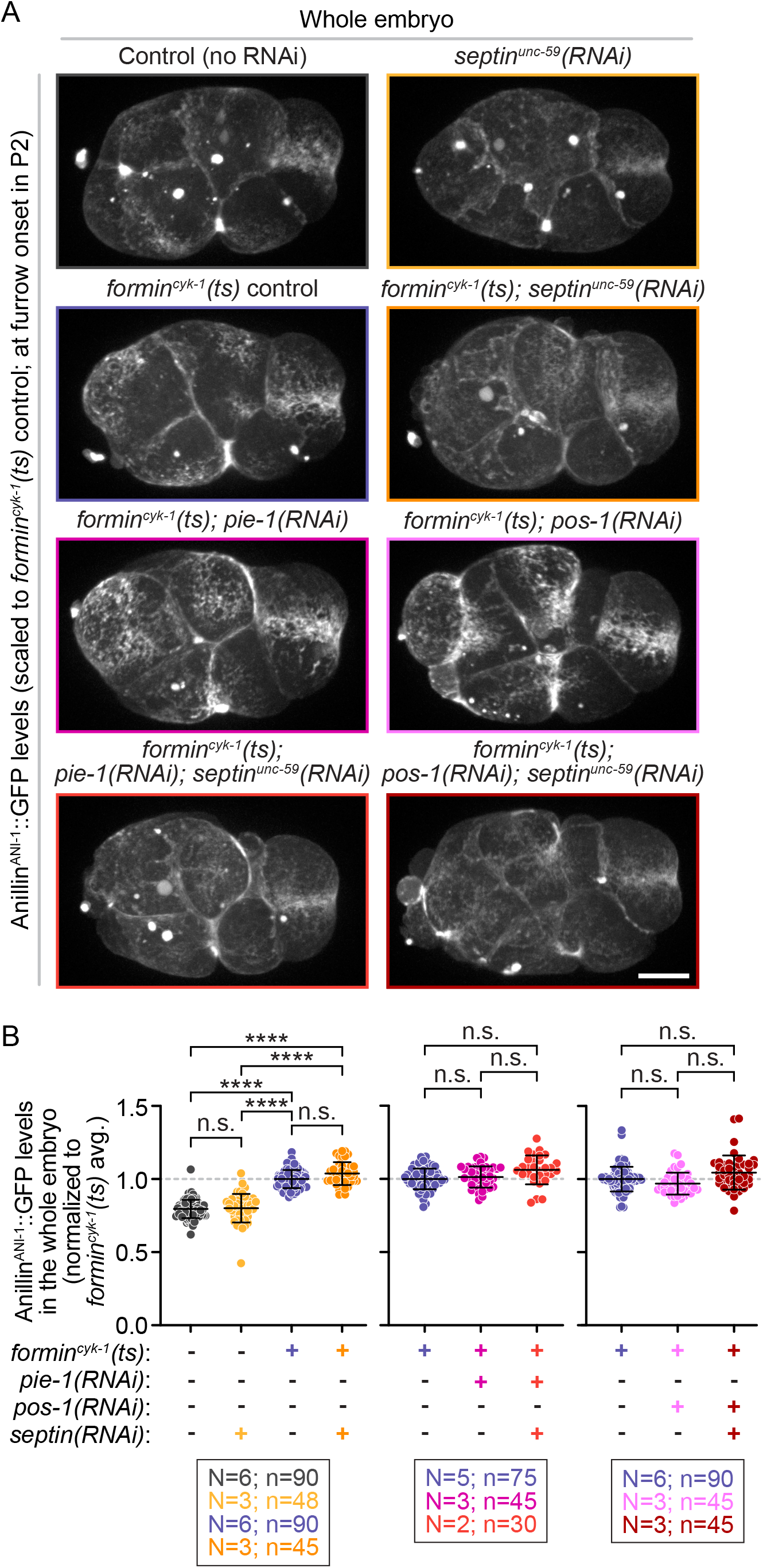
Septin^UNC-59^ depletion does not change anillin^ANI-1^ whole 4-to-8-cell embryo levels. **A)** Representative maximum projection images of endogenously-tagged anillin^ANI-1^::GFP at the time of cleavage furrow onset in P2 for indicated genotypes (images scaled relative to *formin^cyk-1^(ts)* control embryos); scale bar=10 μm (see schematic in **Figure S7F**, see also Methods). **B)** Graphs showing normalized (to *formin^cyk-1^(ts)* control embryos) whole embryo anillin^ANI-1^::GFP levels in the 4-to-8-cell embryo for each genotype. Error bars=standard deviation; N=number of experimental replicates; n=number of embryos scored for each genotype by color; n.s.=not significant, ****=p-value ≤0.0001 (two-way ANOVA, see also **Table S1**).

**Figure S10:**
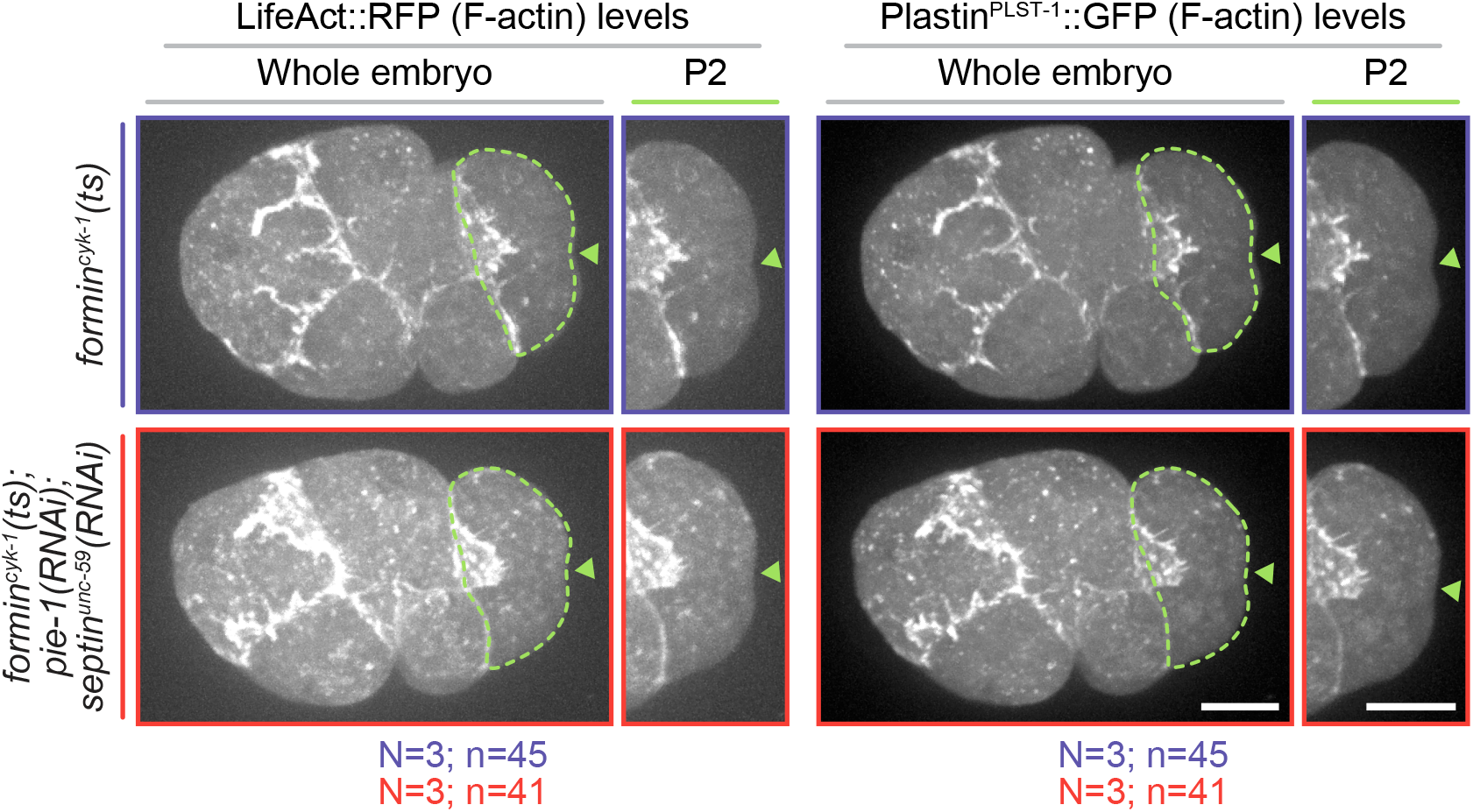
The P2 cell divides without detectable F-actin levels at the division plane in formin^cyk-1^(ts); pie-1(RNAi); septin^unc-59^(RNAi) embryos. Representative maximum projection images of the whole embryo (left images) and P2 cell (right images) in 4-to-8-cell stage embryos expressing the fluorescently tagged F-actin reporters Lifeact::RFP (left panels) and plastin^PLST-1^::GFP (right panels) showing no detectable linear F-actin in the division plane at the time of cleavage furrow onset in P2. scale bar=10 μm; N=number of experimental replicates; n=number of embryos scored for each genotype by color.

